# Population receptive field shapes in early visual cortex are nearly circular

**DOI:** 10.1101/2020.11.05.370445

**Authors:** Garikoitz Lerma-Usabiaga, Jonathan Winawer, Brian A. Wandell

## Abstract

The visual field region where a stimulus evokes a neural response is called the receptive field (RF). Analytical tools combined with functional MRI can estimate the receptive field of the population of neurons within a voxel. Circular population RF (pRF) methods accurately specify the central position of the pRF and provide some information about the spatial extent (diameter) of the receptive field. A number of investigators developed methods to further estimate the shape of the pRF, for example whether the shape is more circular or elliptical. There is a report that there are many pRFs with highly elliptical pRFs in early visual cortex (V1-V3; Silson et al., 2018). Large aspect ratios (>2) are difficult to reconcile with the spatial scale of orientation columns or visual field map properties in early visual cortex. We started to replicate the experiments and found that the software used in the publication does not accurately estimate RF shape: it produces elliptical fits to circular ground-truth data. We analyzed an independent data set with a different software package that was validated over a specific range of measurement conditions, to show that in early visual cortex the aspect ratios are less than 2. Furthermore, current empirical and theoretical methods do not have enough precision to discriminate ellipses with aspect ratios of 1.5 from circles. Through simulation we identify methods for improving sensitivity that may estimate ellipses with smaller aspect ratios. The results we present are quantitatively consistent with prior assessments using other methodologies.

**Significance Statement:** We evaluated whether the shape of many population receptive fields in early visual cortex is elliptical and differs substantially from circular. We evaluated two tools for estimating elliptical models of the pRF; one tool was valid over the measured compliance range. Using the validated tool, we found no evidence that confidently rejects circular fits to the pRF in visual field maps V1, V2 and V3. The new measurements and analyses are consistent with prior theoretical and experimental assessments in the literature.

## Introduction

Small regions of the primate visual cortex (V1-V3) contain neurons whose spatial receptive fields are compact and often overlap in the visual field. The receptive fields of individual neurons can be measured from electrical activity (Hubel and Wiesel, 1968). Using fMRI responses it is possible to measure the receptive fields of individual cortical voxels (Dumoulin and Wandell, 2008). These fMRI responses reflect the activity of many (∼10^5^) neurons and are called the population receptive field (pRF). There has been extensive work using pRF methods to measure visual cortex in the living human brain (Wandell and Winawer, 2015) and versions of these methods with intrinsic and calcium imaging have been used in animal model systems (Kalatsky and Stryker, 2003; Nauhaus et al., 2016).

The population RF estimates depend upon models of the physiological response. The early pRF models used simple linear models of the physiological response, often assuming that the pRF has a canonical spatial profile (e.g., circularly symmetric Gaussian; Dumoulin and Wandell, 2008). Such simple models can accurately predict the fMRI time series of voxels in early visual cortex (e.g., V1-V3) when using a limited range of stimuli, capturing a very large proportion of the explainable variance. Over time investigators have expanded the scope of the stimuli and this required increasing the complexity of the pRF models (Zuiderbaan et al., 2012; Kay et al., 2013; Greene et al., 2014; Alvarez et al., 2015).

This journal published a provocative claim that prior investigators had missed an important aspect of the human population receptive field shapes in V1-V3 (Silson et al., 2018). Nearly all the prior work in which a parametric form was assumed treated the pRF spatial profile as approximately circular. This question had been tested, for example, by Zeidman et al. (2018), who found that elliptical fits are not better than circular models. On the other hand, Silson et al. (2018) report that pRFs are significantly elongated, often with an aspect ratio (ratio of long to short axis) of 2.5 or greater. Groups using a broader class of allowable shapes have reported inconclusive results, for example finding most pRFs in V1-V3 to be nearly circular but a small percentage to be quite elongated (Greene et al., 2014), or finding many voxels to be slightly elongated (Merkel et al., 2018, 2020).

We set out to investigate the discrepancy between the high ellipticity report and the more common assumption of near circularity by replicating the findings. We began the replication by using the same software as in Silson et al. (2018). As part of our workflow we ran the software through a recently developed validation framework (Lerma-Usabiaga et al., 2020). This assessment revealed that the software returns inaccurate estimates, including ellipses with aspect ratios larger than 2 when tested with ground-truth circular data (aspect ratio of 1). To pursue the key scientific question, we decided to use a different software tool and to perform a full assessment of how accurately this tool might measure deviations from circularity. The validation of the second software tool identified a range of conditions where performance is reliable. Using the validated software with retinotopy data from the 7T Human Connectome Project, we find no support for a shape that is substantially and systematically different from circular.

## Materials and Methods

For software validation we used synthetic data generated using the validation framework (pRF-synthesis) described by Lerma-Usabiaga et al. (2020). These methods are described in detail in that paper, and summarized briefly here. We added two levels of noise (low- and mid-noise) created with realistic models of several noise sources, including physiological noise (cardiac and respiratory), low frequency drift, and instrumental noise (white noise) derived from experimental measurements. We used bars with contrast patterns that swept the 20 degree diameter visual field vertically, horizontally and in 45 and −45 degrees, in two different directions each (8 bar sweeps in total). The total stimulus duration and TR (sampling rate) were varied across several simulations. See Table 1 for details.

**Table 1.** Main parameters of the experiments

| Type | Simulated or real TR (sec) | Duration (sec) | Noise | Aspect ratio | Eccentricity (deg) | pRF Size (deg) | pRF-Analysis* |
| --- | --- | --- | --- | --- | --- | --- | --- |
| Synthetic | 2 | 400 | None | {1,2} | {1,2,3,4,5,6,7,8,9} | {.5, 1, 1.5, 2, 3, 4} | {AFNI6, Vista6} |
|  | 2 | 400 | {low, mid} | {1,2} | {1,2,3,4,5,6,7,8,9} | {0.25:0.25:6} | {AFNI6, Vista6} |
|  | 1 | {300,400} | {low, mid} | {1,2} | {1,2,3,4,5,6,7,8,9} | {0.25:0.25:6} | {Vista6} |
| Experimental (V1,V2,V3) | 1 | 300 | NA | NA | Limited to: [2,6] | Limited to: [1,3] | {Vista6, Vista4} |

\* Note: 6 refers to the elliptical fits with 6 parameters, and 4 to the circular fits with 4 parameters.

For assessing deviations from circularity of population receptive fields, we used empirical measurements from the 7T HCP Retinotopy project (Benson et al., 2018). Specifically, we selected retinotopy data collected from the three representative subjects analyzed in Figure 7 of that paper (HCP IDs 164131, 115017, and 536647). We analyzed the empirical data using the containers (pRF-analyze) described, implemented and shared by Lerma-Usabiaga et al., (2020).

### Experimental design and statistical analyses

We evaluated the mrVista (https://github.com/vistalab/vistasoft) and AFNI (Cox, 1996) estimates of elliptical population receptive fields. The latter was introduced in (Silson et al., 2018). The former is part of the mrVista toolbox but has not previously been used in published work.

In mid-2018 the AFNI development team discovered an error in the ellipse formula. Silson et al. (2018) re-ran the analyses and reported some numerical differences but no changes to the pattern of results or the conclusions: the pRF solutions remained highly elongated after correcting the code. Here, we used the new, corrected version of the software. The April 4th 2020 version is implemented in the Docker container. The noiseless analyses in Figures 1 and 2 were performed in a local macOS binary installation with an August 28th 2018 version of Afni. We validated the corrected algorithms using synthetic (ground-truth) input data and estimated the following parameters.

**Fig 1.**
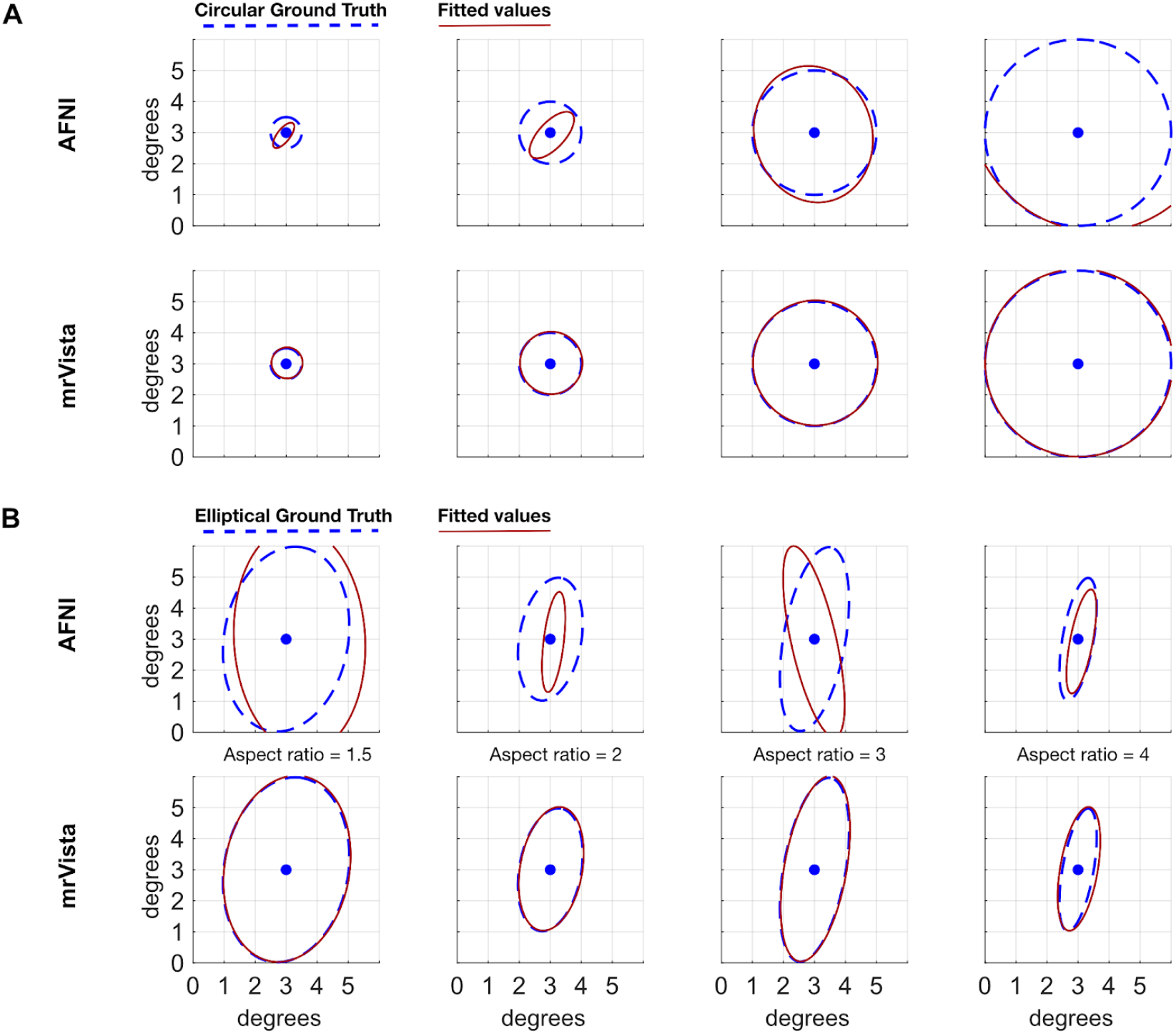
AFNI-elliptical does not recover accurate parameters of noise-free synthetic data. We analyzed noise-free synthetic data analysis with AFNI-elliptical and mrVista-elliptical. (A) AFNI-elliptical (top row) and mrVista-elliptical (bottom row) analyses of circular, Gaussian, ground-truth data with four different pRF sizes. The dashed line represents the 1 SD radius of the Gaussian. (B) Same as A but with elliptical ground truth data.

**Fig 2.**
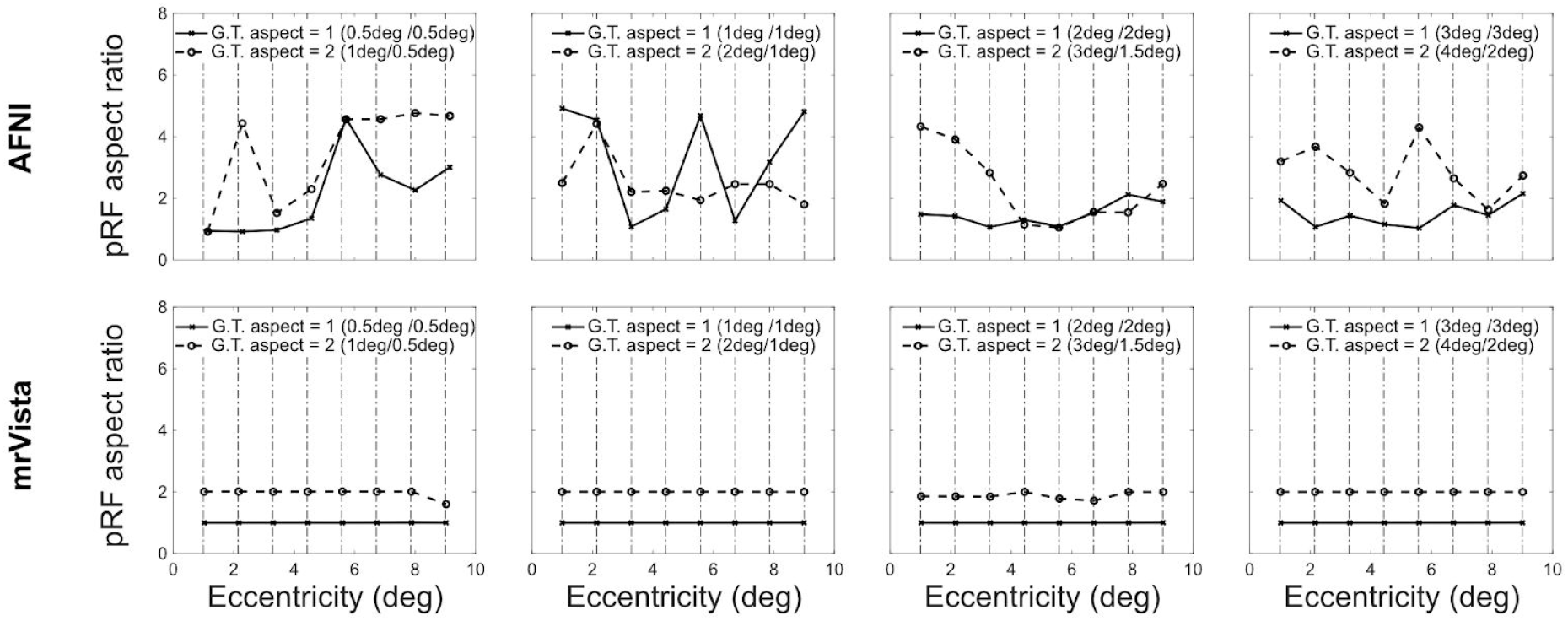
AFNI-elliptical has systematic aspect ratio errors at different eccentricities and sizes. AFNI-elliptical (top row) and mrVista (bottom row) results for circular (aspect ratio 1, solid lines) and elliptical (aspect ratio 2, dashed lines) ground truth synthetic time series, with pRF radii ranging from 0.5 deg to 4 deg. *G*.*T*.: *Ground Truth*.

- The center position of the population receptive field (x, y).
- The standard deviations (*σ*_1_, *σ*_2_) of the two axes of the ellipse (*σ* 1> *σ*_2_). (Circular fits are constrained to *σ*_1_ = *σ*_2_ and one parameter is returned.)
- The angle *θ* of the main axis (larger sigma). Not returned for the circular fit.
- The gain parameter *A*.

We estimated deviations from circular pRFs using the mrVista prf-Analyze container and measurements obtained from the HCP project. The pRF-Analyze-mrVista container returns the distribution of aspect ratio estimates (*σ*_1_,/*σ*_2_). We compared median values and distributions from fitting the empirical measurements with values expected from analyzing ground-truth data generated using prf-Synthesize.

Several analyses were performed in this manuscript using synthetic or real data with the following parameters.

### Code availability and reproducibility

To reproduce the computations in this paper requires that Matlab and Docker be installed on your computer. The configuration files and the HCP data for the empirical analyses are curated and stored in a project at the Open Science Foundation (https://osf.io/9jhcm/). The software we describe downloads the data from that OSF project.

The code specific to this paper is shared in the GitHub repository PRFmodel that is within the vistalab project (https://github.com/vistalab/PRFmodel.git). After cloning that repository, please select the git tag EllipsePaperv02. Place this repository on your Matlab path. The script pmMainEllipseFiguresScript.m describes how to install the necessary support libraries and execute the relevant scripts.

The software for the pRF-Validation framework (Lerma-Usabiaga et al., 2020), including the code used to synthesize the BOLD time series, is shared in the same repository. The Docker container image can be downloaded from Docker hub with the command *docker pull garikoitz/prfsynth*. The mrVista analysis code is publicly shared in github.com/vistalab/vistasoft, and its container can be downloaded from Docker hub with the command *docker pull garikoitz/prfanalyze-vista*. The AFNI analysis code is publicly shared in github.com/afni/afni, and our containerized version used for the analyses in this paper can be downloaded from Docker hub with the command *docker pull garikoitz/prfanalyze-afni*.

## Results

We present results about algorithm validity in noise-free and simulated noise conditions. We then define a range of parameters in which one algorithm performs acceptably, and we analyze empirical measurements from that range. In previous work we reported that pRF algorithms systematically misestimate pRF parameters if there is a mismatch between the hemodynamic response function (HRF) used to simulate the time series with the HRF assumed in the analysis tool. Throughout the simulations here, we used synthetic data that matched the expected HRF.

### Algorithm validity: noise-free analyses

We first set out to validate elliptical models in AFNI (“AFNI-elliptical”) and mrVista (“mrVista-elliptical”) using noise-free synthetic data. We synthesized the BOLD time series for pRFs that are circular and centered at (3,3) deg, with radii spanning 0.5 − 3 deg. For these conditions, AFNI-elliptical inaccurately estimates the pRFs as elongated rather than circular, whereas mrVista-elliptical estimates nearly circular pRFs. We then validated the two algorithms with elliptical ground truth data (Figure 1B). The ground-truth pRFs were again centered at (3,3) deg, and had aspect ratios between 1.5 and 4. Again, AFNI-elliptical fails to estimate the parameters accurately and mrVista-elliptical succeeds.

To explore whether there are systematic errors in AFNI-elliptical or mrVista-elliptical, we synthesized a noise-free dataset by systematically varying eccentricity, size and aspect ratio (Figure 2). The AFNI-elliptical algorithm generally returns incorrect aspect ratios. Over these parameter ranges, the mrVista-elliptical algorithm generally returns accurate estimates.

### Algorithm robustness: noise analyses

In the presence of measurement noise, the aspect ratio of circular pRFs will be overestimated. Suppose that the ground truth is a circle with radius r. The major and minor axes will both be estimates of the true radius plus noise,*r* + *Ñ*. The estimated aspect ratio, A, is the ratio of the two noisy samples constrained

so that the major axis is the larger of the two samples

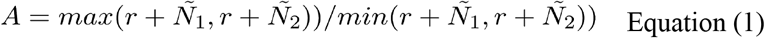

From this formula we observe that (1) the estimated value must be greater than 1, and (2) the impact of the noise will be large when the pRF radius is small. We performed numerical simulations of the formula in Equation 1, using a range of radii and plausible noise distributions (Supplementary Figure S6). We observed that for small radii starting at 0.25 deg the median aspect ratio is of 2.5. The median aspect ratio values reduce asymptotically towards the aspect ratio of 1 as the radius increases.

We tested AFNI-elliptical and mrVista-elliptical using simulated noisy datasets for a circular pRF with a radius of 2 deg (Figure 3). The simulated stimulus had a TR=2, bar width of 2.8 deg and step size of 1.2 deg; the simulated duration was 400 seconds, including 8 bars sweeps across the visual field. The time series were identical for 100 simulations except for different random samples of noise. The AFNI-elliptical algorithm estimates the center location accurately, but it does not estimate the aspect ratio as expected. There are many large aspect ratios (> 4), and there are many estimates of 1.0 which should be rare given the noise. The mrVista-elliptical algorithm estimates the center location accurately. The median aspect ratio is generally in the range between 1.2-1.4, as expected. AFNI requires the aspect ratios to be bounded, so we restricted them to be between 1 and 5. MrVista has no such requirements and therefore the aspect ratios were unbounded.

**Fig 3.**
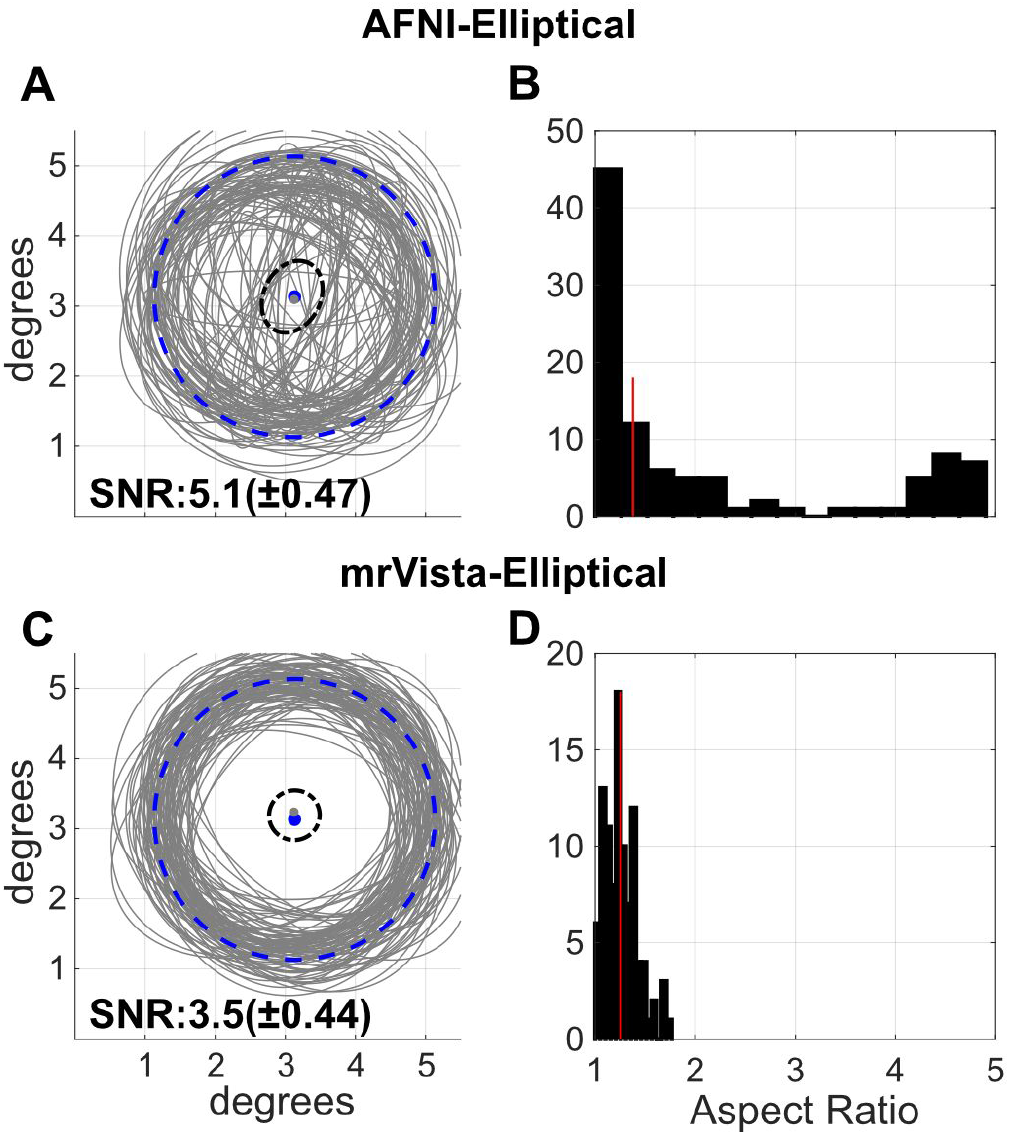
AFNI-elliptical estimates include large aspect ratios for circular ground-truth data with added noise. AFNI elliptical (top row) and mrVista (bottom row) analysis results for 100 noisy simulations (low noise). On the left, the representation of all the receptive fields (gray) over the ground truth (blue dashed line); the black dashed line contains the center locations and the blue dashed line represents the 1 SD radius of the Gaussian. On the right, the histogram of the aspect ratios. The median is indicated by the red line. SNR is the mean and STD of all 100 bold time series. Due to differences in the HRFs between the two algorithms and randomization used in the synthesis, the average SNR of the simulated time series differs, being lower for mrVista.

Based on the simulations, we expect the estimated aspect ratio of circular, noisy ground-truth data to be slightly larger than 1. The mrVista-elliptical estimates conform to this expectation: they are distributed compactly around an aspect ratio of 1.26 ± 0.16 (Figure 3D). The AFNI-elliptical estimates (Figure 3B) are very different, with a larger mean and a much larger standard deviation: 2.07 ± 1.36. Critically, AFNI returns many aspect ratio estimates greater than 3 which suggests that such values should not be taken as evidence of large aspect ratios in the data. A paired t-test comparing the magnitude of the aspect ratios showed that AFNI-elliptical’s error is significantly bigger than mrVista-elliptical’s (t: 6.7, p: 1.4e-09).

To test the generality of these findings, we synthesized and analyzed ground truth datasets with a broader range of parameters. For AFNI-elliptical (Figure 4), the estimated aspect ratio for circular pRFs was about 2.5-3.0 for all ground truth radii (Figure 4A). The distribution of values is quite wide, spanning all the aspect ratios within the 1,5 bounds (Figure 4B). When the ground-truth aspect ratio was 2, the estimated aspect ratio increased to a median value of 4, but the distribution remained very broad. These simulations used the mid-level of noise (see Methods). The results were similar for the low-level noise (see Figure S5a), and various eccentricity values (see Figures S3-S4). These validation tests reveal that, with our configuration, environment variables and function calls, AFNI-elliptical is not a suitable tool for assessing the aspect ratio of population receptive fields. The validation tests produce a wide range of aspect ratio distributions, as found in the empirical analyses reported by Silson et al. (2018). The reason for the differences in these AFNI distributions is unknown.

**Fig 4.**
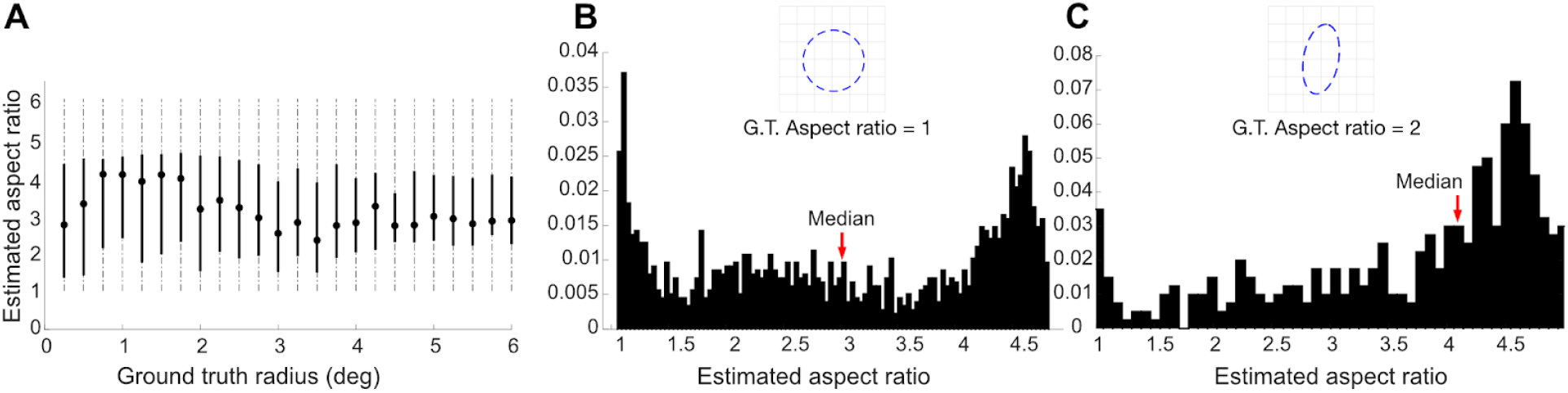
AFNI-elliptical does not estimate the correct aspect ratio of synthetic data. (A) Estimated aspect ratio (ground truth aspect ratio = 1) as a function of pRF radius (deg). The points are the median and the lines show the range corresponding to the central 50% of the estimates. (B) Histogram of estimated aspect ratios (ground truth aspect ratio = 1) using simulated pRFs with a mixture of radius sizes (1-4 deg) and eccentricities (2-6 deg). (C) Histogram of estimated aspect ratios (ground truth aspect ratio = 2) for the same mixture of radius sizes and eccentricities. The simulated bar width is 2.8 deg and the bar translates 1.2 deg for each TR (2 sec). The simulations used the mid-level of noise. The red arrow indicates the median value of the histogram. *G*.*T*.: *Ground Truth*.

We next analyzed the ability of mrVista-elliptical to accurately estimate the pRF aspect ratio (Figure 5). As expected from the basic analysis of signal to noise (Equation 1), the accuracy of the aspect ratio estimates depends on pRF size and properties of the stimulus. With a TR of 2 s, simulations with small pRF radius (∼1 deg or less) are very inaccurate, including many large aspect ratios (Figure 5A). We simulated the accuracy of recovering a circular pRF using a mixture of pRF sizes (1-4 deg) and eccentricities (2-6 deg). The median estimated aspect ratio is approximately 1.5, with the estimates falling mostly between 1 and 2 (Figure 5B). Simulating with a ground-truth aspect ratio of 2 increases the median, but the estimates are spread over a large range (Figure 5C).

**Fig 5.**
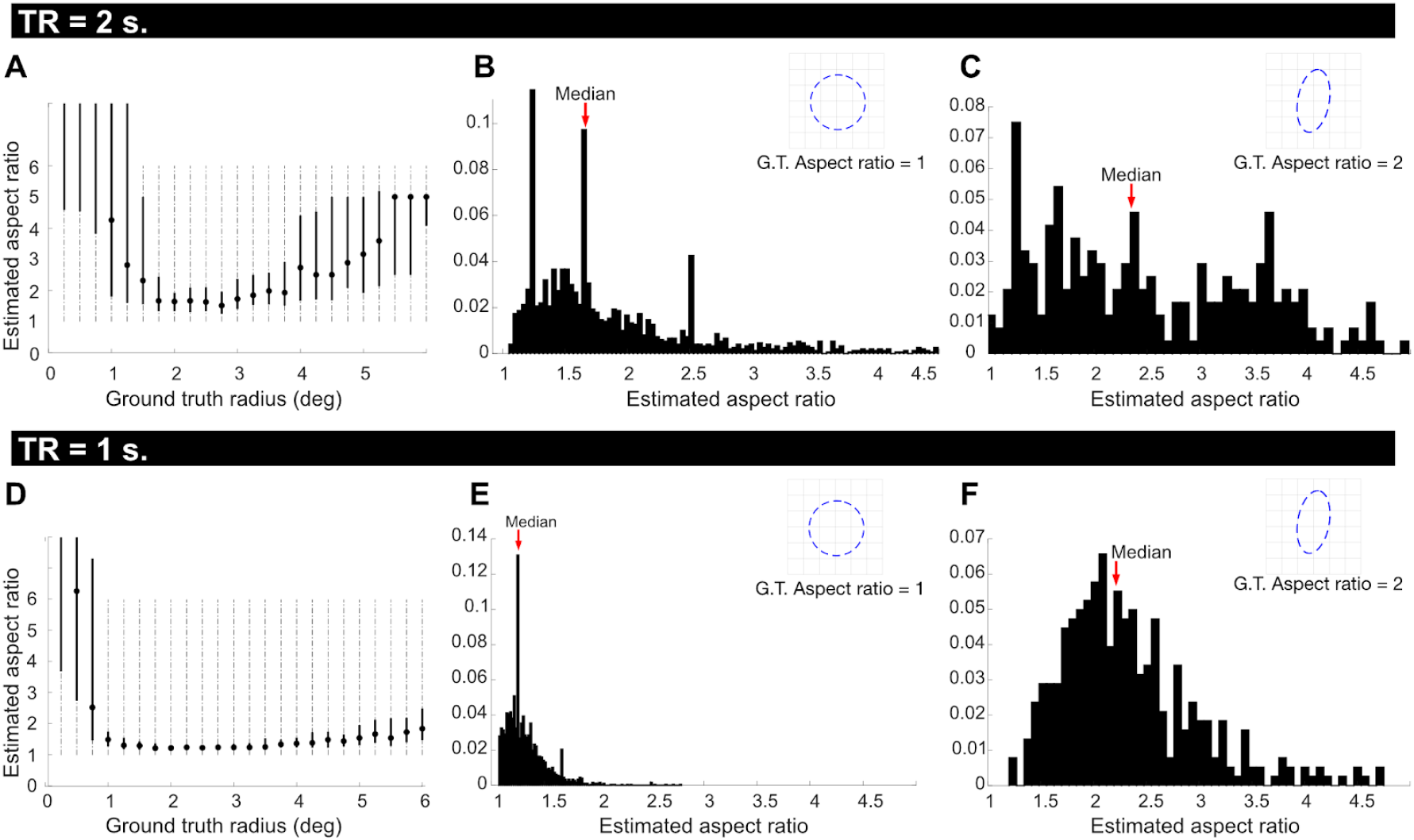
mrVista-elliptical aspect ratio estimates are close to accurate over a limited range of conditions. (A) Estimated aspect ratio (ground truth = 1) as a function of pRF radius (deg). The points are the median and the lines show the range corresponding to 50% of the estimates. (B) Histogram of estimated aspect ratios (ground truth = 1) using simulated pRFs with a mixture of radius sizes (1-4 deg) and eccentricities (2-6 deg). (C) Histogram of estimated aspect ratios (ground truth = 2) for the same mixture of radius sizes and eccentricities. The simulated bar width is 2.8 deg and the bar translates 1.2 deg for each TR (2 sec). The simulations used the mid-level of noise. (D-F) The same graphs calculated with a smaller bar displacement (0.6 deg) and shorter TR (1 sec). For the large bar step size the mrVista elliptical estimates differ between the ground-truth aspect ratios of 1 and 2, although there is very poor accuracy when the ground truth aspect ratio is 2. Reducing the bar step size and increasing the number of temporal samples improves the accuracy of the aspect ratio estimate (D-F). The peaks in the histogram at aspect ratios of 1.25, 1.6 and 2.5 in B, E are a flaw in the algorithm. These peaks, which are present in fits to empirical data (below), are likely due to the coarse-to-fine search method implemented in the algorithm. These simulations define a range of experimental parameters where mrVista-elliptical provides useful information about the aspect ratio. *G*.*T*.: *Ground Truth*.

To understand how empirical methods using mrVista might impact algorithm validity, we carried out simulations with different experimental parameters. Specifically, we simulated an experimental protocol with a shorter TR (1 s instead of 2 s), corresponding to a smaller stimulus step size. The mrVista-elliptical estimates are more accurate over a larger range (Figure 5D). Estimating ground-truth pRFs with a range of sizes and eccentricities, the median aspect ratio is 1.25 and the range is more compact (Figure 5E). The estimates for a ground truth aspect ratio of 2 and multiple pRF sizes have a median aspect ratio slightly larger than 2 (2.2). The same analysis performed with a 2-s TR and low noise results in estimates similar to the 1-s TR simulations (see Figure S5a).

The mrVista-elliptical algorithm has a numerical estimation error that biases the results to return certain aspect ratios (1.25, 1.65, 2.5); these are the peaks in the histogram. We suspect this failure arises from the multi-resolution (coarse to fine) search methodology. The coarse fit uses a grid of parameter values, and the values of the aspect ratio in the grid include these three values. This limitation of the algorithm does not render it unusable for further exploration with real measurements under certain conditions.

### Empirical measurement: estimated aspect ratio

We used mrVista-elliptical and experimental data to assess the aspect ratio of pRFs in early visual cortex. We analyzed three typical subjects from the HCP 7T retinotopy dataset (Benson et al., 2018). We first used mrVista-circular to assess the parameter ranges. The estimated range of eccentricity values (2.5-6.5 degs) and the pRF areas (6.5-30 deg^2^) were then used to restrict the mrVista-elliptical fits. These parameters are consistent with many previous estimates and place no restriction on the estimated aspect ratio values. The HCP dataset was acquired with a TR of 1 s and a duration of 300 secs, and our analysis of synthetic data above indicates that in this spatial step, eccentricity and size range, the median mrVista-elliptical aspect ratios are reliable. Each subject had two runs with sweeping bar stimuli, and we analyzed the average of these runs.

We also created synthetic datasets with low- and mid-noise levels to compare with the experimental data (Figure 6). The synthetic datasets used the same sequence of stimulus apertures as the experimental data.

**Fig 6.**
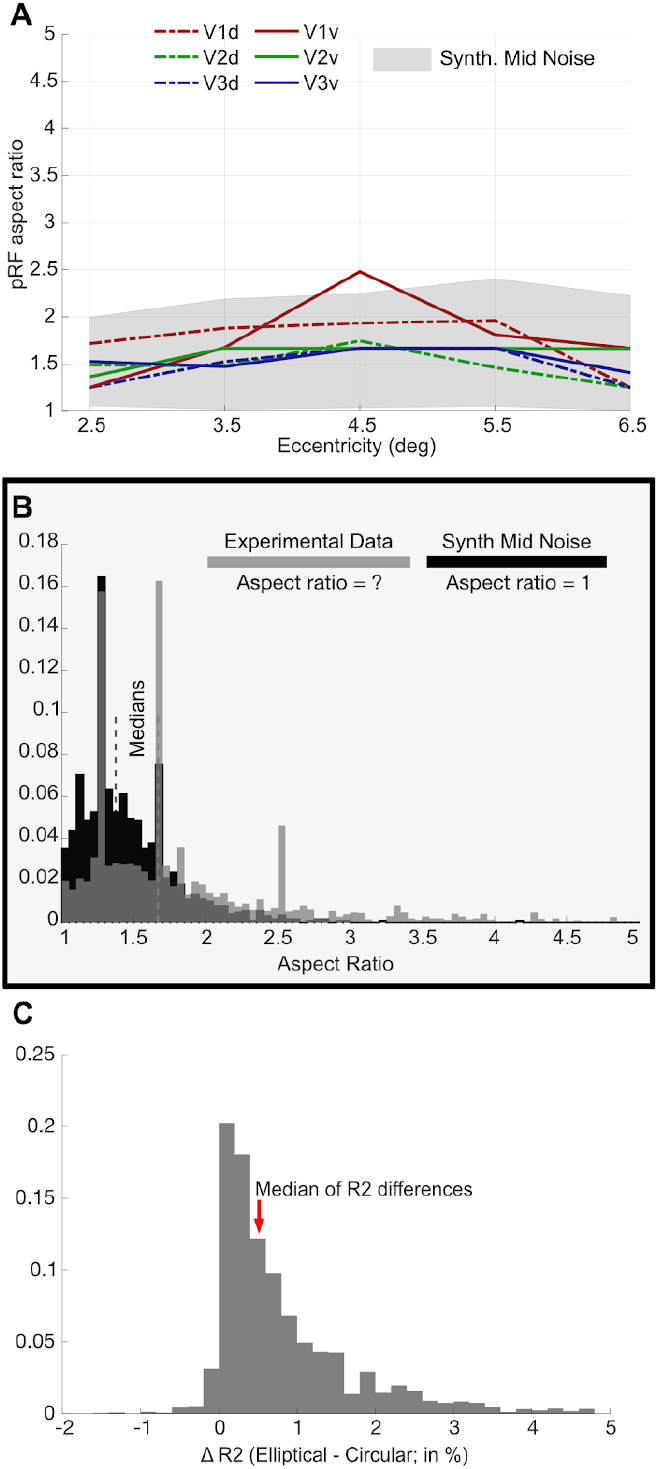
mrVista-elliptical pRF parameters estimated from empirical measurements in V1-3 (N=3). (A) Estimated median pRF aspect ratios of experimental (color) data plotted as a function of eccentricity. The experimental data are plotted separately for ventral and dorsal regions of V1-3. Synthetic data were created using mid-level noise and are represented as a light gray band containing the central 95% aspect ratio values. The experimental data aspect-ratio fits show a large variance across voxels, but except one case, the population medians are within the expected range of the (mid) noise simulations. (B) Histograms of the estimated pRF aspect ratio for experimental (gray) and synthetic (black) data. Estimates were included in the histograms if the model fit explained at least 25% of the variance and the pRF position was between 2.5-6.5 deg and the pRF area size estimate was between 6.5-30 deg^2^. The ground truth aspect ratio for the synthetic data was 1. The thin dashed vertical lines represent the median values for the experimental and synthetic analyses. (C) Histogram of the difference in variance explained (R^2^) between the elliptical and circular model fits to the experimental data. The histogram includes data subject to the same restrictions as in (B). The precise parameters for determining the restriction do not impact the conclusions in either (B) or (C).

**Fig 7.**
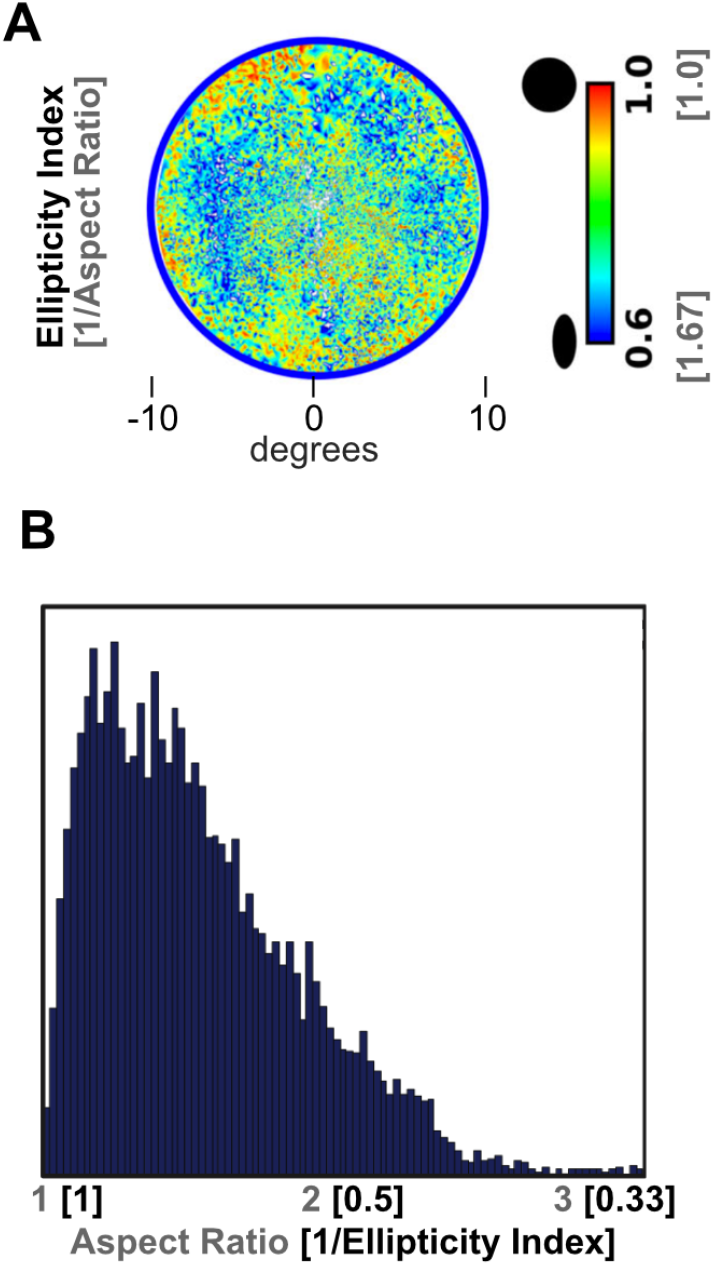
Ellipticity results reported in the literature show similar results to our circular ground truth simulations. (A) Ellipticity (1/Aspect Ratio) reported in (Merkel et al., 2020). (B) Aspect ratio reported in (Greene et al., 2014).

The analyses of the experimental data returned a median pRF aspect ratio of about 1.5, with no systematic effects of ventral vs dorsal, or eccentricity (Figure 6A). The values are slightly larger in V1. There is no significant trend in the aspect ratio as a function of eccentricity. The aspect ratio from the experimental data is similar to the ratio in the mid-noise synthetic data. The experimental data aspect-ratios have higher variance than the synthetic values, but they are within the 95% confidence interval of the mid-noise synthetic aspect ratio estimations. As expected, the low-noise synthetic data simulations have aspect ratios closer to the ground truth value. This indicates that the mid-noise level of the synthetic data is a good approximation to the experimental data (Figure 6B). When comparing experimental data from all maps, the histograms are similar, with median aspect ratio values at around 1.5. The validation procedures (Figure 5) show that mrVista can accurately capture aspect ratios of 2, and yet the empirical data return a distribution of aspect ratios that is smaller, closer to the theoretically expected values of a circular ground truth.

Finally, we analyzed the strength of the evidence in favor of using an elliptical model compared to the circular model. The elliptical model uses two more parameters and contains the circular model as a special case. Hence, it is expected that the variance explained (R^2^) will be higher for the elliptical model. In rare cases, the R^2^ is lower for the elliptical model, indicating a failure of the optimization to find the best solution. Figure 6C shows the histogram of the difference between the R^2^ of the elliptical and circular fits, for all experimental data in which the models explain more than 25% of the variance. The elliptical fit is systematically higher than the circular fit, but the median difference is less than 1%, and even the few voxels with the largest difference are no more than 5%. Hence, there is almost no evidence in support of using the elliptical model over the circular model for these experimental data. Detecting differences from circularity will require new protocols and models.

Many analyses of these types of histograms, separating the data in various ways such as dorsal and ventral or by visual field map, support the same conclusions (see Figure S7). The median aspect ratio values remain between 1.25 and 1.5, and we found no systematic relationship between the estimated aspect ratio or ellipse orientation and pRF position in the visual field.

## Discussion

### The application of pRF methods

Multiple groups have used population receptive field parameters as dependent variables to understand the effects of cortical plasticity, attention, and diagnostic tools for neurology and psychiatry (Wandell and Winawer, 2015). For example, pRF methods have been used to examine hypotheses about brain substrate changes, such as excitation-inhibition imbalances, that may be associated with neurological, ophthalmologic and psychiatric diseases (Papanikolaou et al., 2014; Wandell and Winawer, 2015; Anderson et al., 2017; Dumoulin and Knapen, 2018). There are also opportunities to understand individual differences in the visual pathways that may impact performance in tasks that rely on vision, such as reading (Le et al., 2017) and face recognition (Witthoft et al., 2016). Establishing the precision of the parameter estimates obtained with current protocols and tools enables us to determine with more confidence whether an individual under study is within the distribution of typical subjects.

### Oriented pRFs

What would be a plausible biological basis for pRFs with large aspect ratios? Many neurons in primary visual cortex have oriented receptive fields. For simple cells, the spatial envelope tends to be elongated along the axis of orientation tuning (De Valois et al., 1982; Ringach, 2002; Michel et al., 2013). The neurons are arranged in an orderly pattern such that the main orientation changes smoothly across the cortical surface. The receptive fields span many orientations over a 1 mm distance (Hubel and Wiesel, 1974). In typical 3T measurements, a single fMRI voxel aggregates the response over a millimeter or more and thus accumulates the metabolic response from neurons with many orientations. It would be quite surprising if the orientation of neuronal receptive fields could be observed robustly in functional MRI measurements. Although there have been some claims to this effect (Kamitani and Tong, 2005; Sasaki et al., 2006; Freeman et al., 2011), it appears now that these biases are due to properties of the stimulus aperture rather than to orientation tuning (Carlson, 2014; Roth et al., 2018)

Alternatively, the pRF from a voxel could be elongated if the neural RFs center positions within a voxel had an asymmetric distribution. Such asymmetric distributions might occur if, for example, the cortical magnification differed systematically between the radial and tangential directions. The effect of neural RF distribution within a voxel on the shape of the pRF is, however, likely to be modest (Amano et al., 2009). Were the pRF measurements truly to have a large aspect ratio, we would still need to find a plausible biological basis.

We are unaware of claims other than Silson et al. (2018) that one can reliably measure a large aspect ratio based on the fMRI response from individual voxels. Direct comparisons of standard pRF models suggest that circular receptive field models provide the best fits (Zeidman et al., 2018; Figure 10). Using novel measurement approaches, investigators report that individual fMRI voxels may have some orientation preference with a magnitude similar to the values reported here (Greene et al., 2014; Merkel et al., 2018, 2020).

For example, Merkel et al. (2018; 2020) estimated the aspect ratios of voxels in early visual cortex and reported ellipticity (the inverse of aspect ratio) ranging between 0.6 and 1, which corresponds to aspect ratios of 1 to 1.67 (Figure 7A; Merkel et al., 2020). Using tomographic methods to estimate pRF shapes (Greene et al., 2014) also estimated aspect ratios (Figure 7B). The distribution they report had 11% of the aspect ratios greater than 2 which is close to the expected amount based on our simulations with synthetic data assuming circular pRFs (14%) and based on analysis with the HCP 7T (8%) data.

These aspect ratios are not meaningfully different from circular given the expected level of experimental noise and current protocols. Specifically, by definition the estimated aspect ratio value must exceed 1. Further, the impact of experimental noise will be quite large when the pRF radius is small. For example, using 0.7 deg standard deviations as noise, for small radii (0.25 − 1 deg) the expected median aspect ratio is almost 3 (see Supplementary Figure 6). This value reduces asymptotically towards the aspect ratio of 1 for large pRF sizes (> 5 deg). Moreover, pRF size estimates are less robust than pRF center estimates and the absolute value depends strongly on the individual HRFs (Lage-Castellanos et al., 2020; Lerma-Usabiaga et al., 2020).

These principles and simulations show accurate estimation of aspect ratio values as small as 1.5 will require new experimental paradigms that mitigate instrumental noise and account for the computational uncertainties. It would also be preferable to use methods that include an accurate assessment of the individual subject’s HRF. Elsewhere we used simulations to describe adjustments to experimental protocols that should improve the accuracy and stability of pRF measurements (Lerma-Usabiaga et al., 2020). Implementing and validating these methods will require some patience.

## Conclusion

This project began with a report that the aspect ratios of pRFs in early visual cortex are substantially larger than previously thought (Silson et al., 2018). We set out to investigate this report, and we concluded that the difference could be traced to a software implementation. Our new data analysis confirmed the prior consensus about pRF shapes in early visual cortex: the best-fitting shapes are not very different from circular (Greene et al., 2014; Zeidman et al., 2018; Merkel et al., 2020). The ability to measure shapes with greater precision, perhaps revealing systematic deviations at individual voxels or even orientation maps, will require advances in protocols and analyses. Simulations suggest these may be in reach (Figure 5).

The complexity of modern neuroimaging analyses has arrived at a point where explicit and public validation frameworks are important for building trust in publications and as part of the standard for software distribution. Here, we used the validation framework implemented in (Lerma-Usabiaga et al., 2020). The development of validated models and quantified parameter estimates has been a hallmark of sensory science, and we continue that approach here.

## Acknowledgements

This project has received funding from the European Union’s Horizon 2020 research and innovation programme under the Marie Sklodowska-Curie grant agreement No 795807 to G.L.-U. and NIH grants supporting J.W. (EY027401, EY027964, MH111417). While writing this paper we have been in contact with E. Silson, C. Baker, and R. Reynolds. We thank R. Reynolds for help with the AFNI software.

## SUPPLEMENTARY FIGURES

**Fig S1a.**
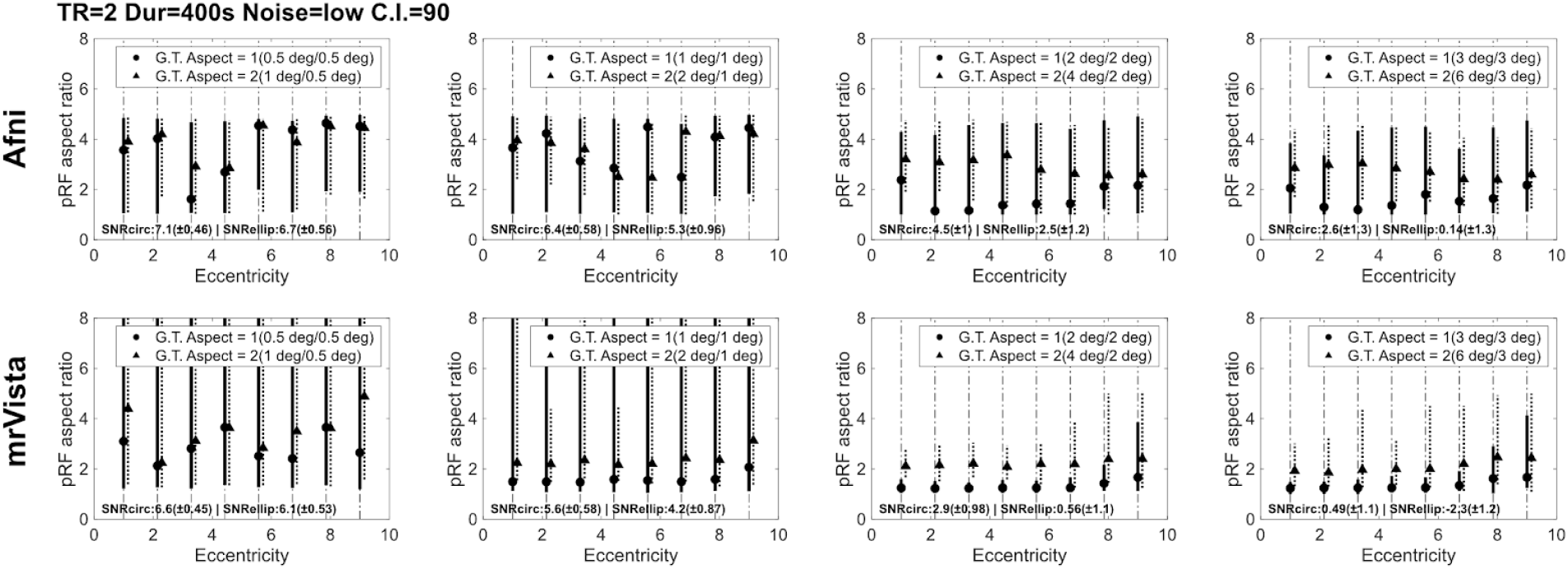
Synthetic data. Eccentricity vs Aspect. AFNI and mrVista. Low noise and 90% CI. TR = 2s, Duration 400s. See main text for the rest of details.

**Fig S1b.**
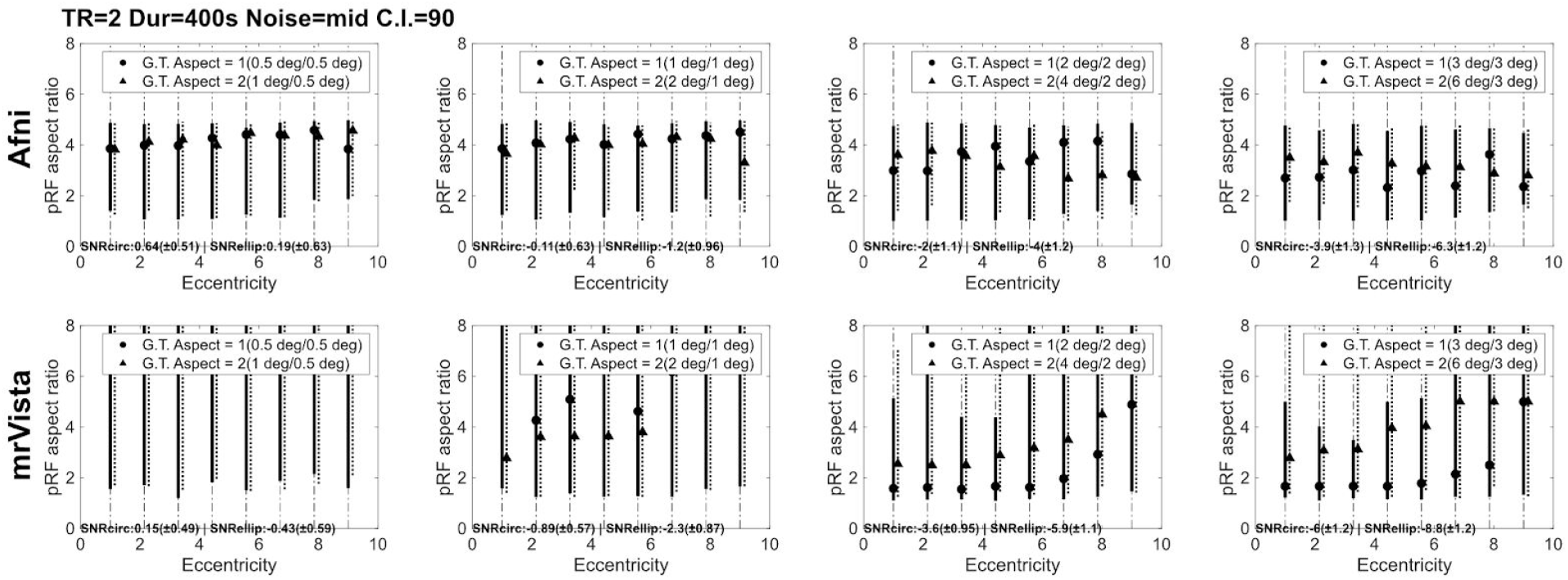
Synthetic data. Eccentricity vs Aspect. AFNI and mrVista. Mid noise and 90% CI. TR = 2s, Duration 400s. See main text for the rest of details.

**Fig S1c.**
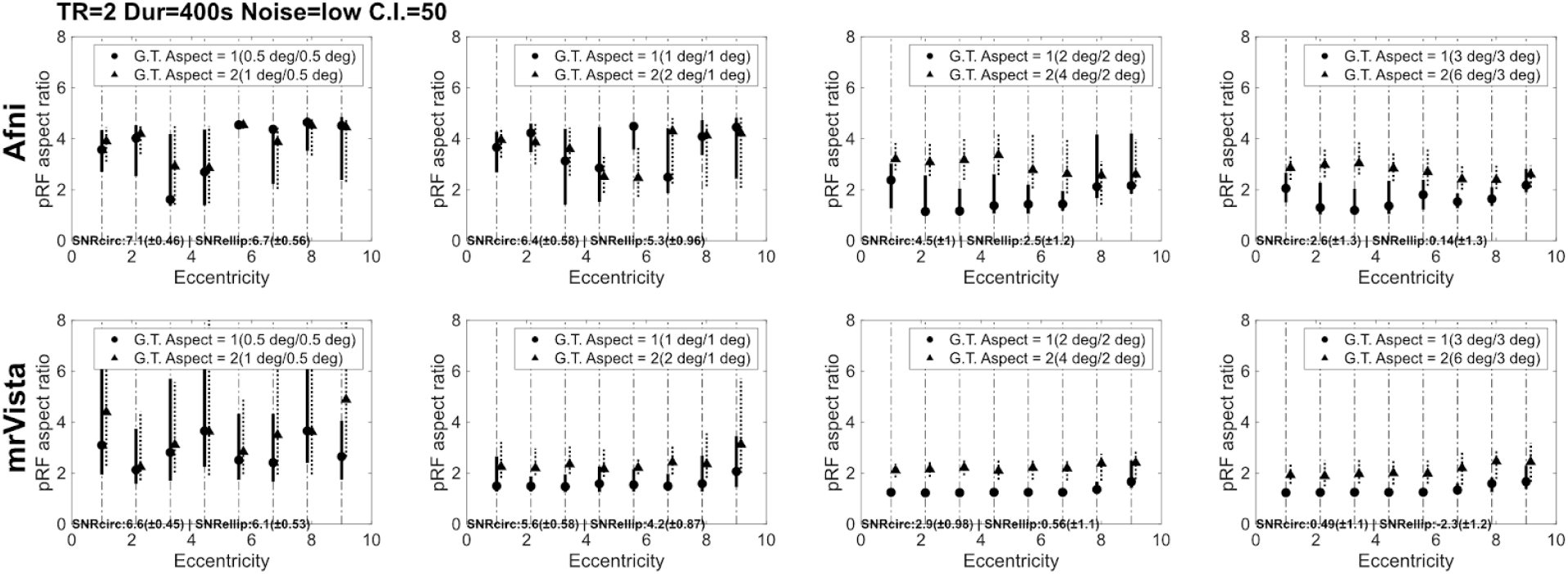
Synthetic data. Eccentricity vs Aspect. AFNI and mrVista. Low noise and 50% CI. TR = 2s, Duration 400s. See main text for the rest of details.

**Fig S1d.**
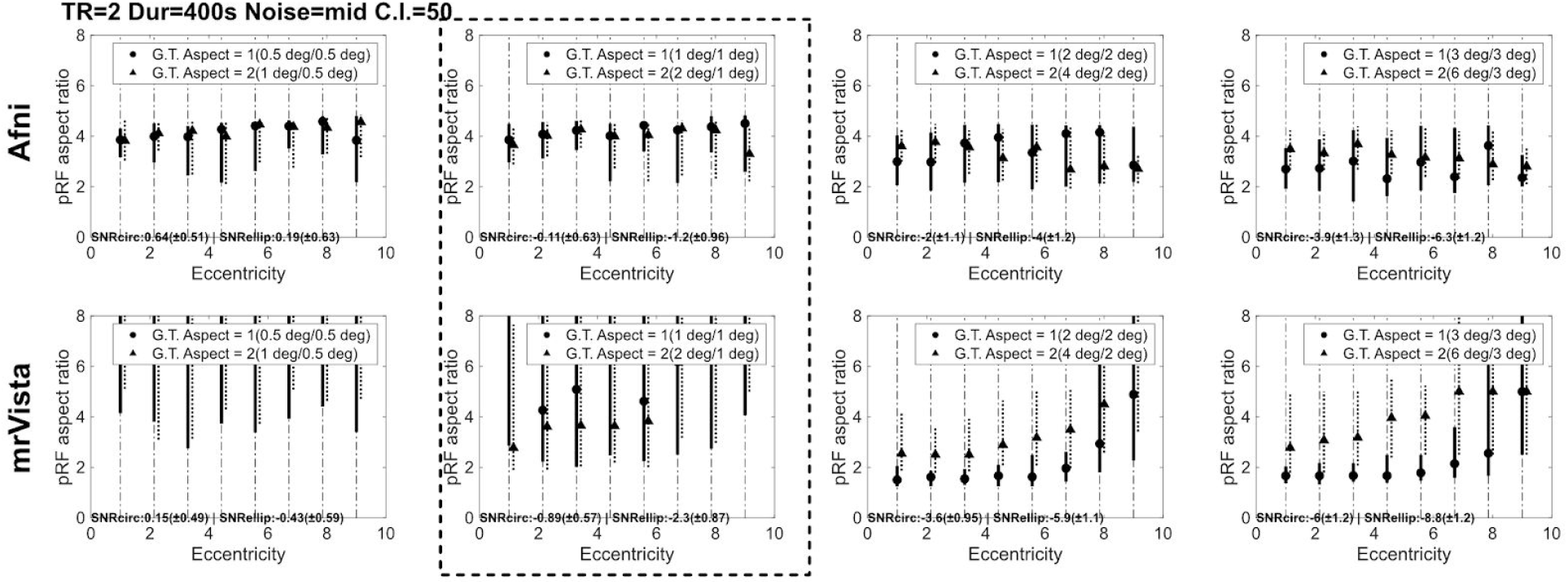
Synthetic data. Eccentricity vs Aspect. AFNI and mrVista. Mid noise and 50% CI. TR = 2s, Duration 400s. See main text for the rest of details. Panes inside a dashed square used in main text.

**Fig S2a.**
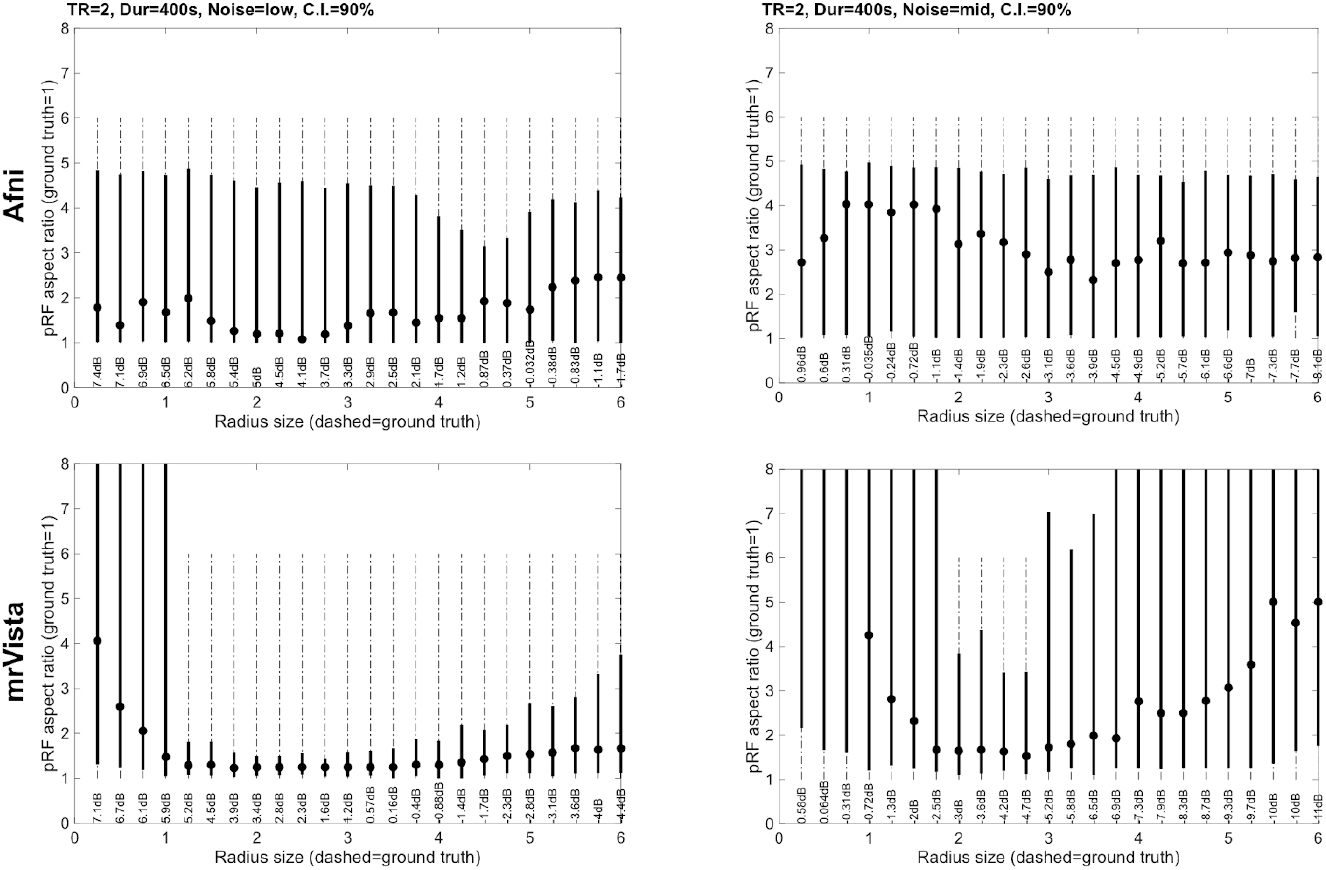
Synthetic data. Size vs Aspect. AFNI and mrVista. 90% CI. TR = 2s, Duration 400s. See main text for the rest of details. Ground truth aspect ratio = 1.

**Fig S2b.**
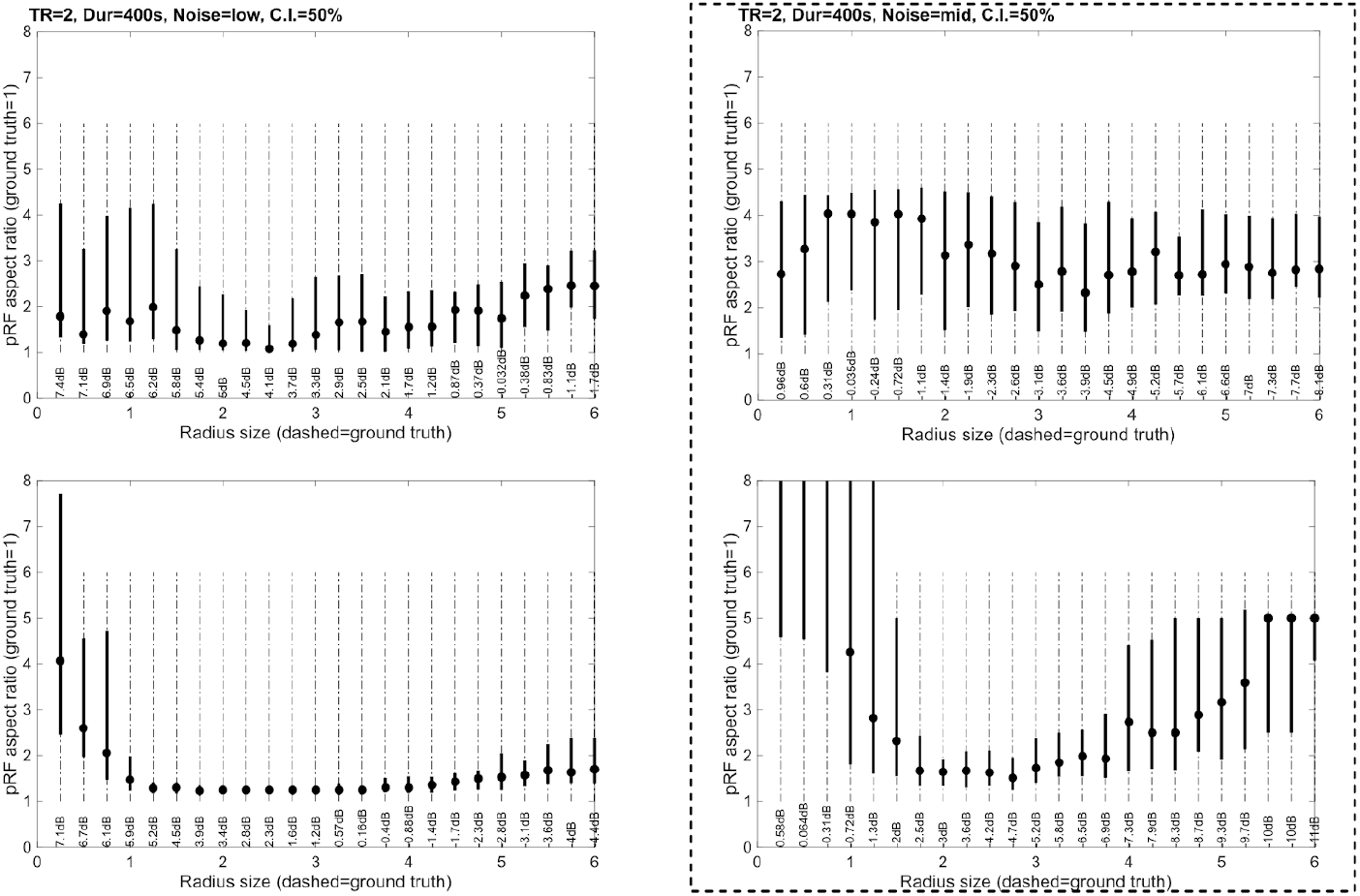
Synthetic data. Size vs Aspect. AFNI and mrVista. 50% CI. TR = 2s, Duration 400s. See main text for the rest of details. . Ground truth aspect ratio = 1. Panes inside a dashed square used in main text.

**Fig S3a.**
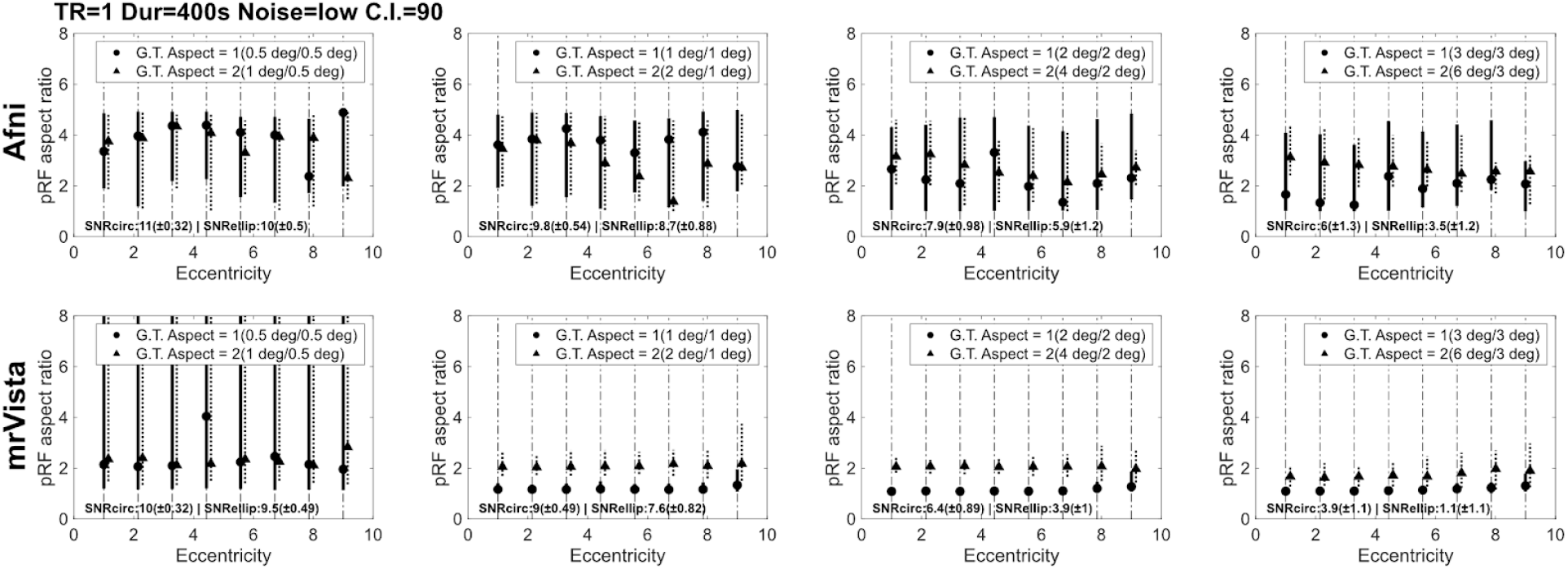
Synthetic data. Eccentricity vs Aspect. AFNI and mrVista. Low noise and 90% CI. TR = 1s, Duration 400s. See main text for the rest of details.

**Fig S3b.**
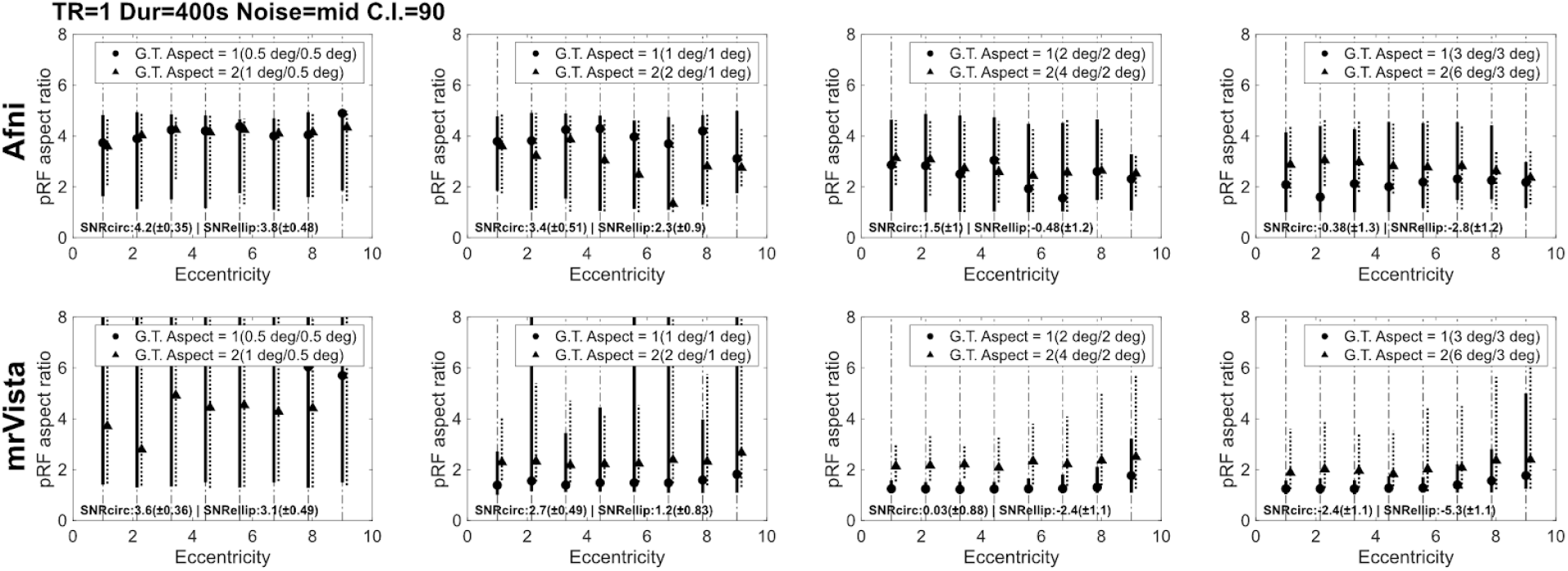
Synthetic data. Eccentricity vs Aspect. AFNI and mrVista. Mid noise and 90% CI. TR = 1s, Duration 400s. See main text for the rest of details.

**Fig S3c.**
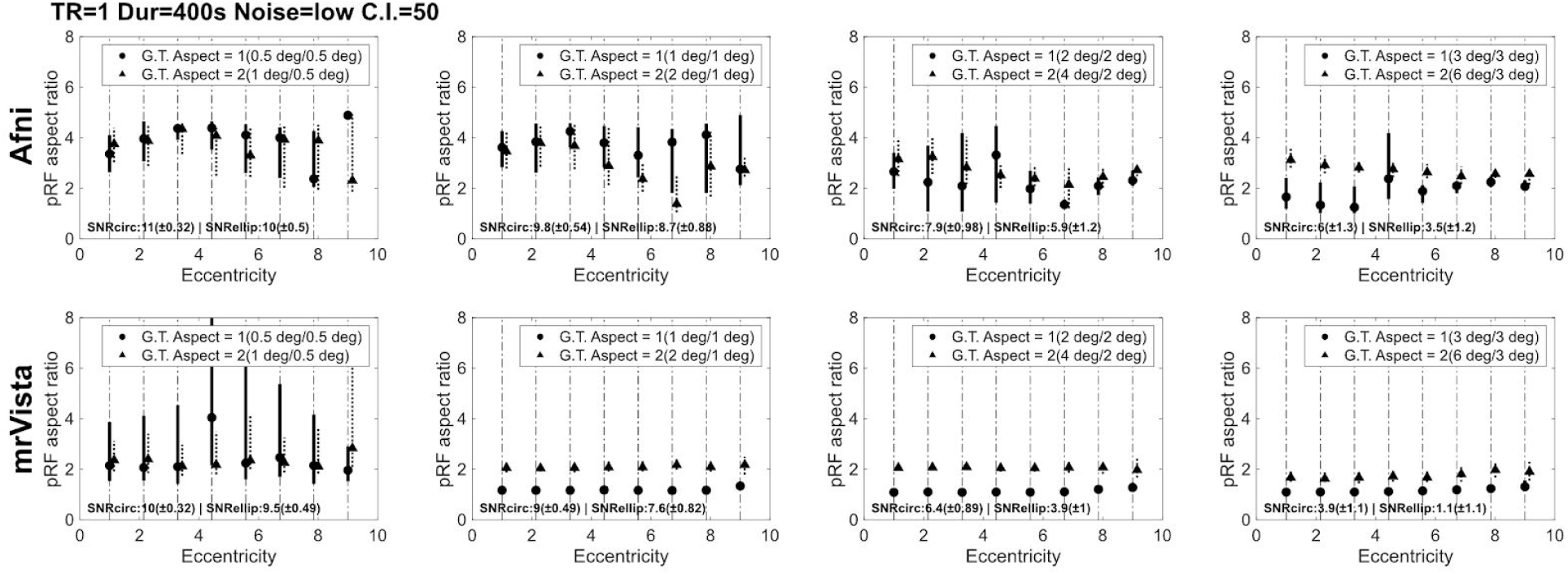
Synthetic data. Eccentricity vs Aspect. AFNI and mrVista. Low noise and 50% CI. TR = 1s, Duration 400s. See main text for the rest of details.

**Fig S3d.**
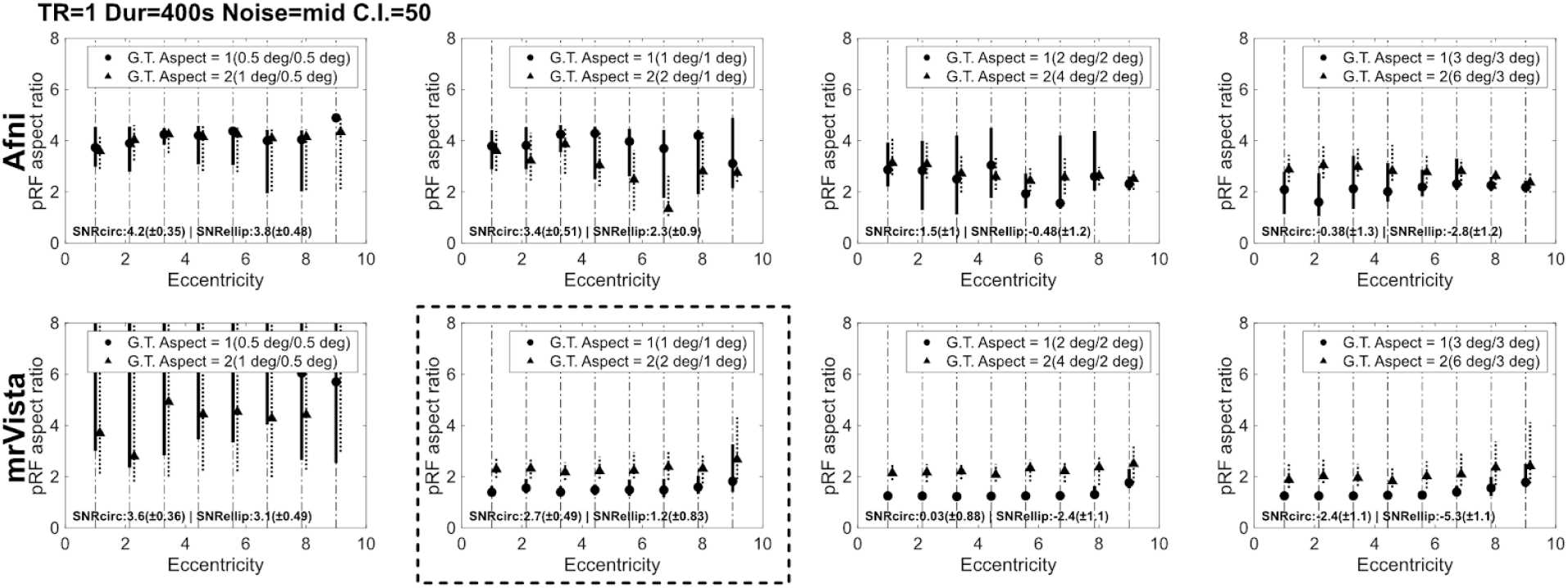
Synthetic data. Eccentricity vs Aspect. AFNI and mrVista. Mid noise and 50% CI. TR = 1s, Duration 400s. See main text for the rest of details. Panes inside a dashed square used in main text.

**Fig S4a.**
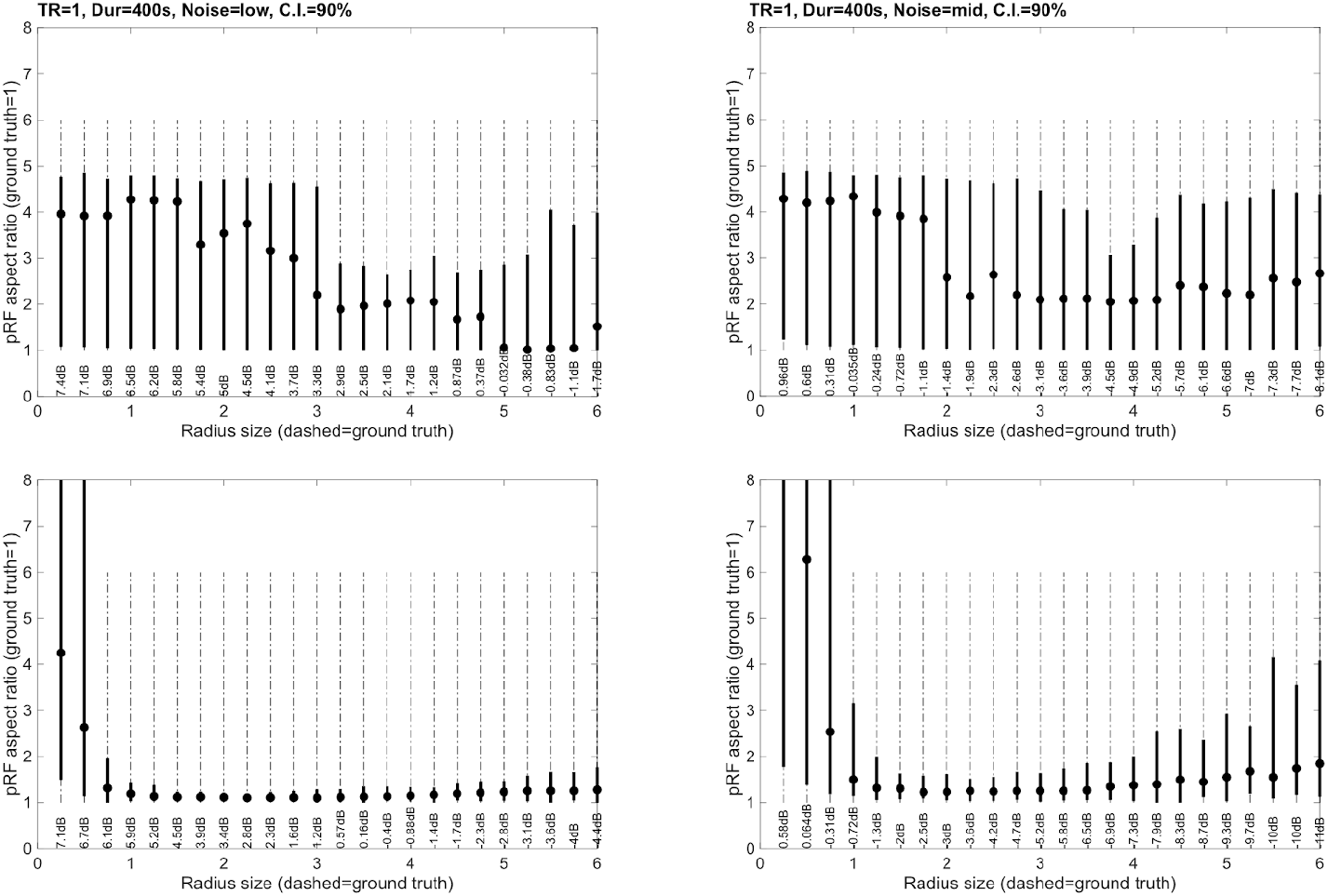
Synthetic data. Size vs Aspect. AFNI and mrVista. 90% CI. TR = 1s, Duration 400s. See main text for the rest of details.

**Fig S4b.**
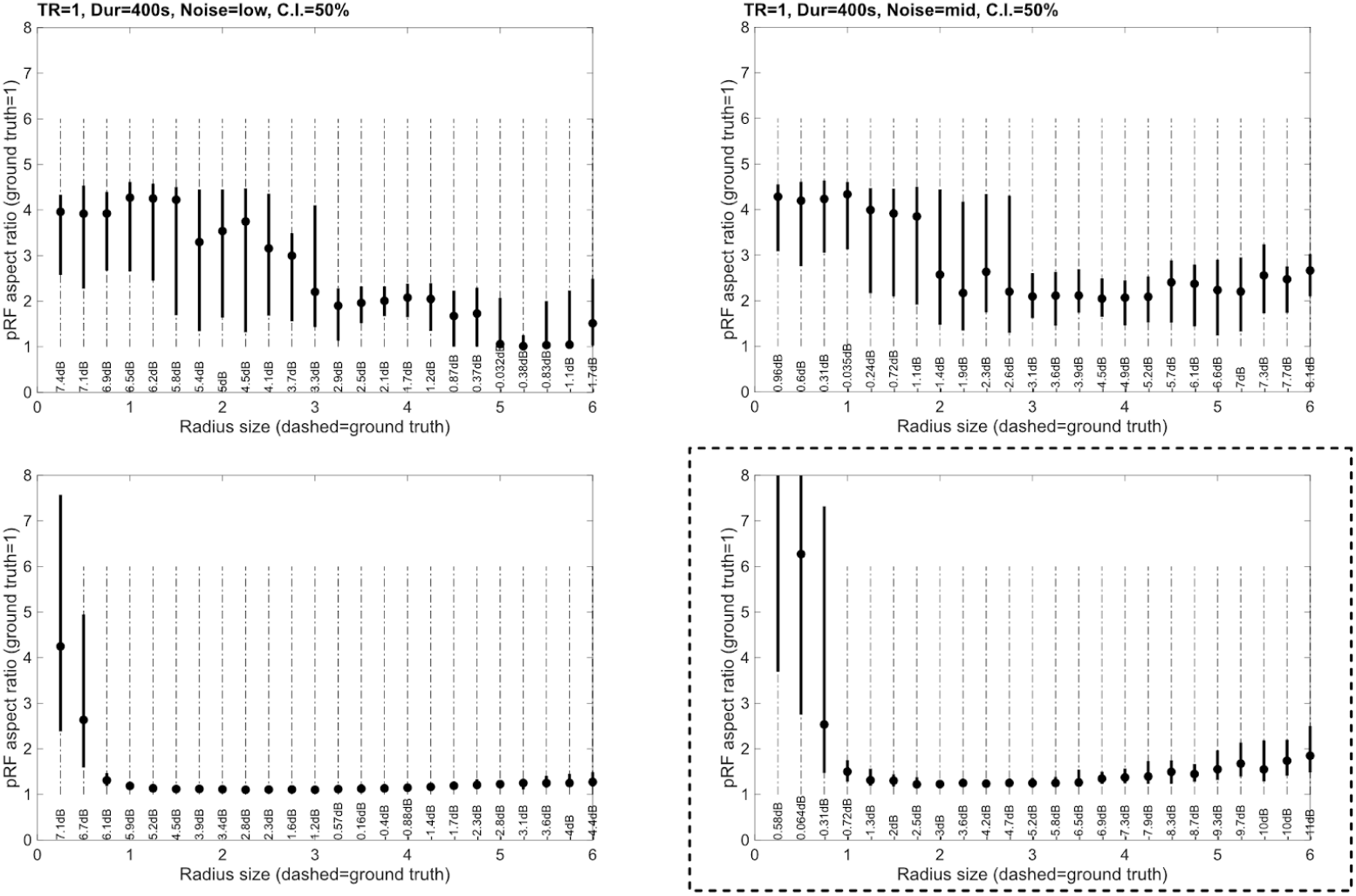
Synthetic data. Size vs Aspect. AFNI and mrVista. 50% CI. TR = 1s, Duration 400s. See main text for the rest of details. Panes inside a dashed square used in main text.

**Fig S5a.**
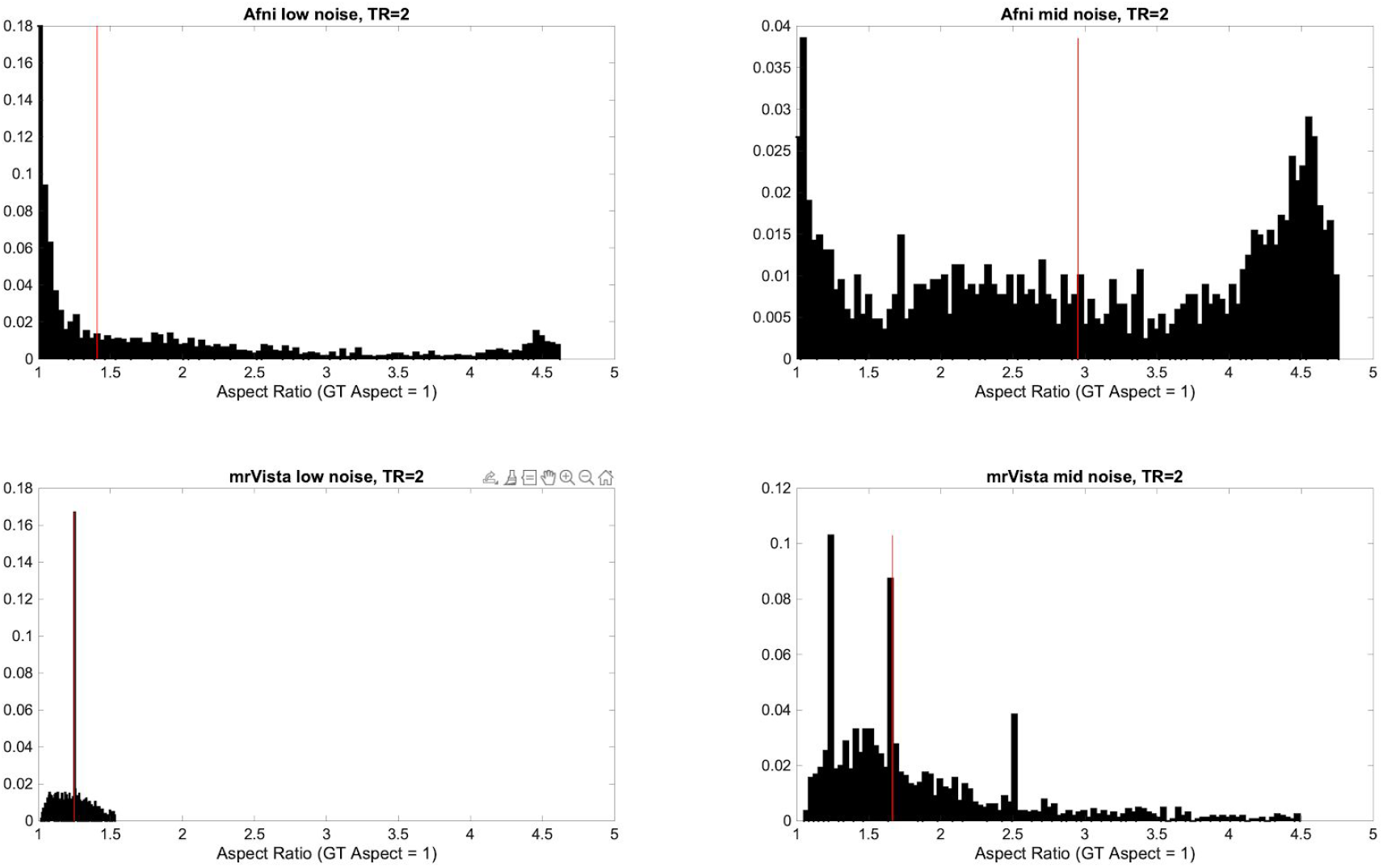
Synthetic data (G.T.= 1). Aspect histograms. AFNI and mrVista. 90% CI. TR = 2s, Duration 400s. See main text for the rest of details. Panes inside a dashed square used in main text.

**Fig S5b.**
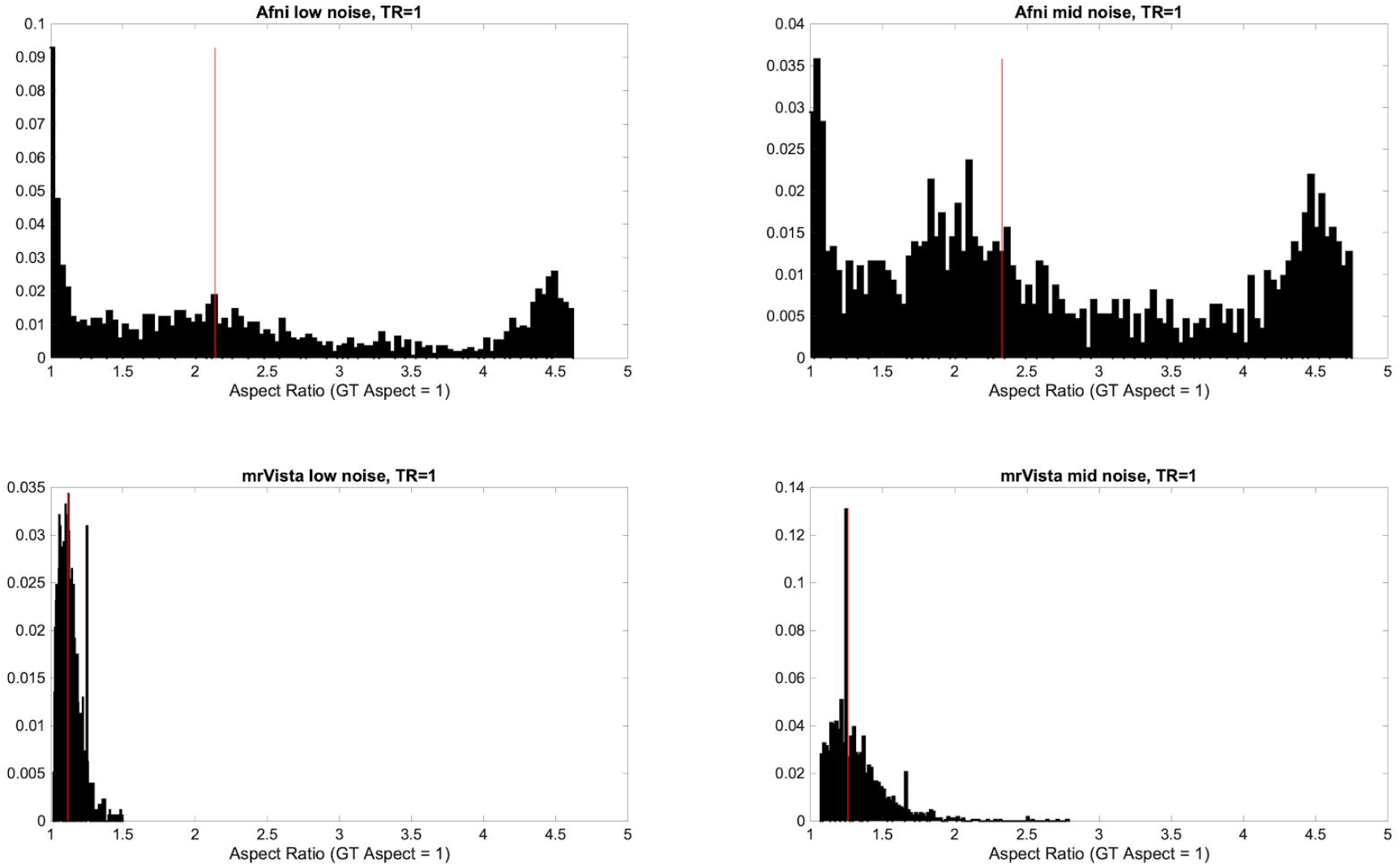
Synthetic data (G.T.= 1). Aspect histograms. AFNI and mrVista. 90% CI. TR = 1s, Duration 400s. See main text for the rest of details. Panes inside a dashed square used in main text.

**Fig S5c.**
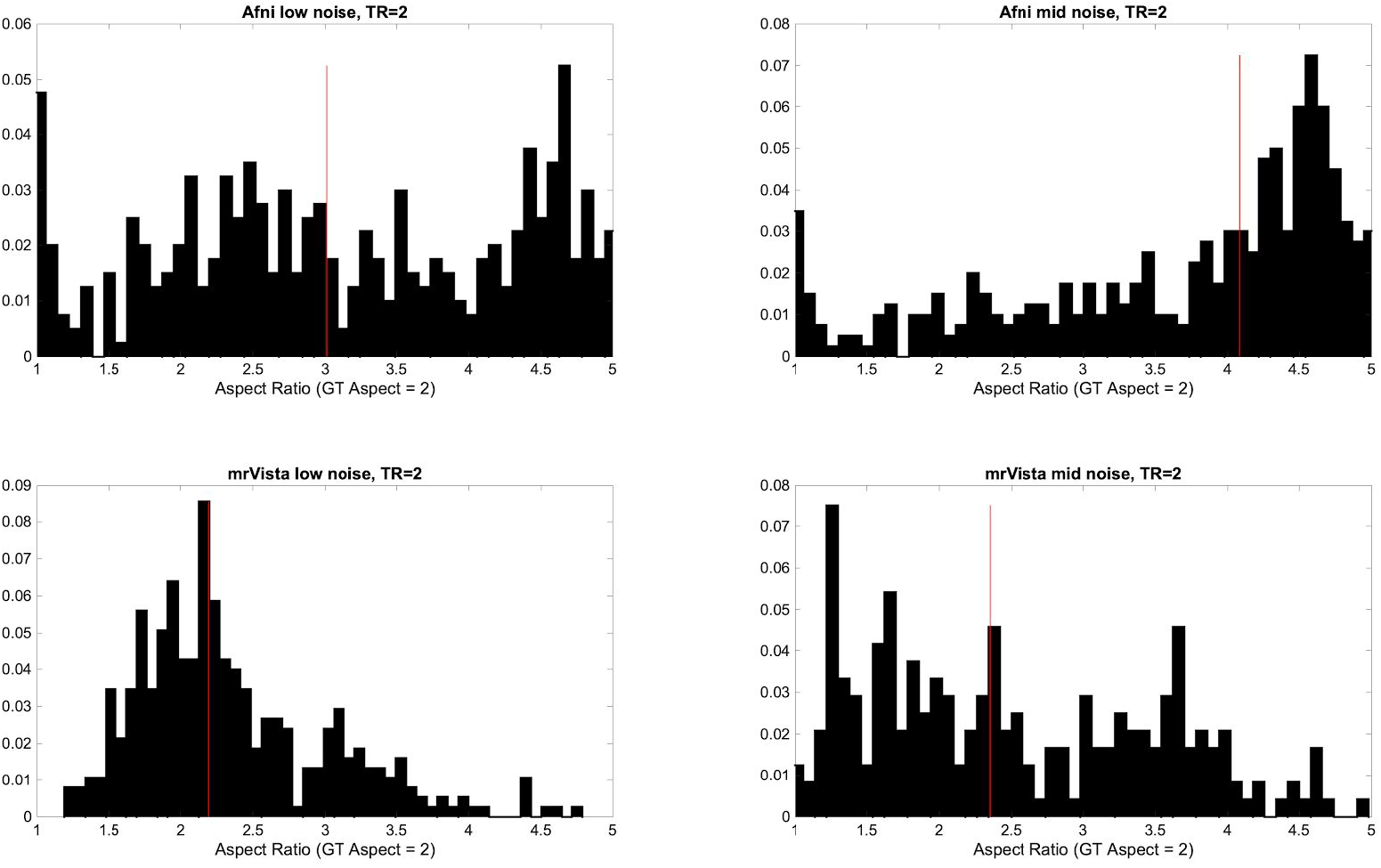
Synthetic data (G.T.= 2). Aspect histograms. AFNI and mrVista. 90% CI. TR = 2s, Duration 400s. See main text for the rest of details. Panes inside a dashed square used in main text.

**Fig S5d.**
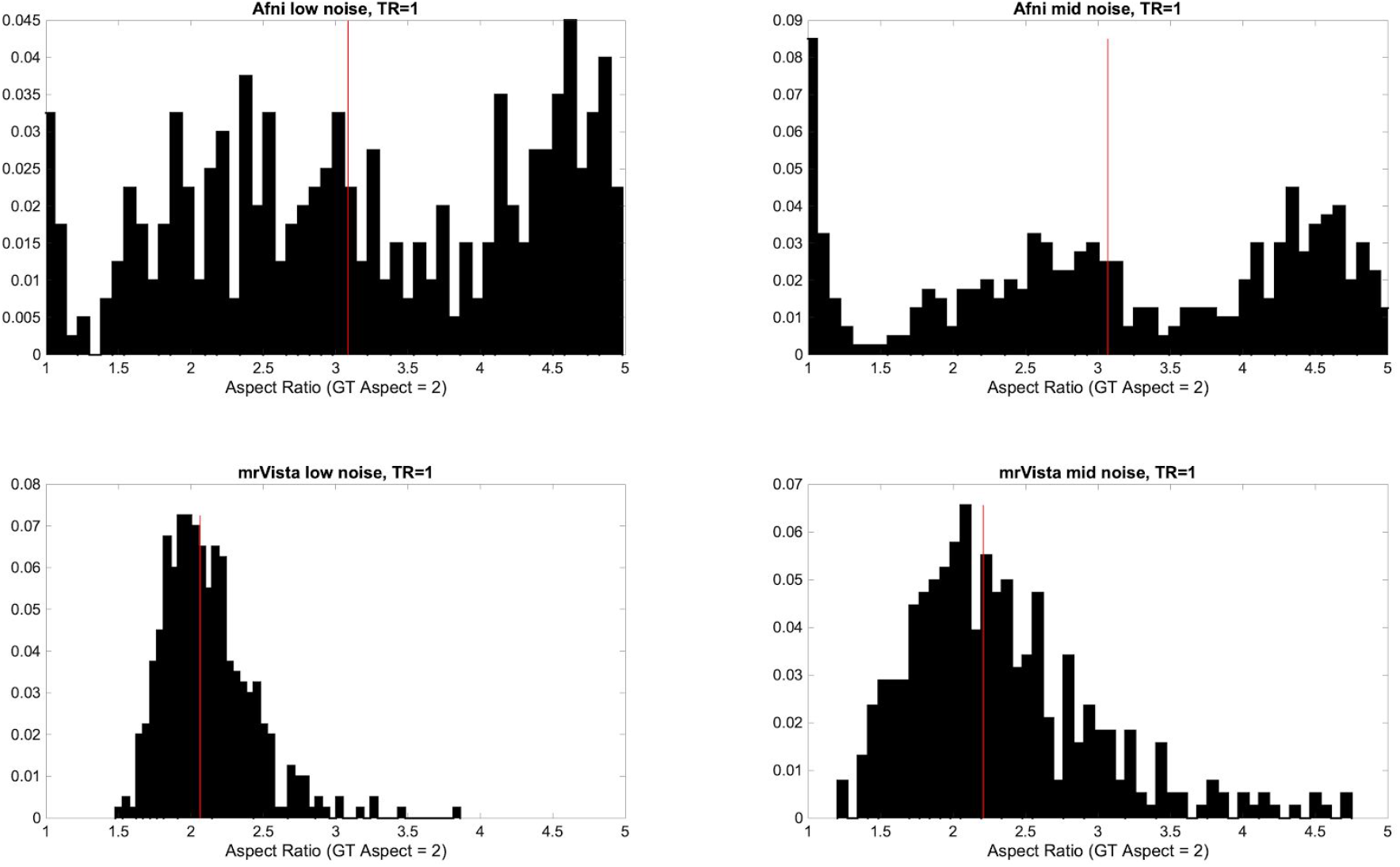
Synthetic data (G.T.= 2). Aspect histograms. AFNI and mrVista. 90% CI. TR = 1s, Duration 400s. See main text for the rest of details. Panes inside a dashed square used in main text.

**Fig S6.**
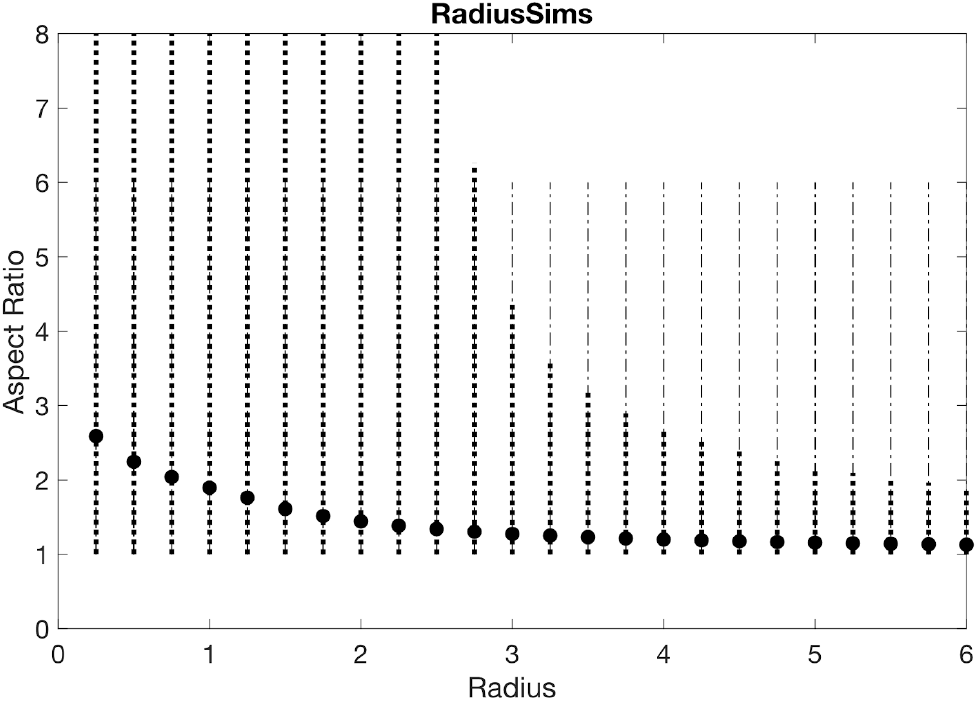
Circle aspect ratio simulations with noise in radius size estimations. r1 = r2 take the values in the x axis, and 1000 random noise values are added (Noise Factor = .7) to obtain R1 and R2. We consider only R1 > 0.01 and R2 > 0.01 at the same time. With the resulting pairs, we calculate the ratio that is always positive with the formula: Aspect Ratio = max([R1,R2],[],2) ./ min([R1,R2],[],2);

**Fig S7.**
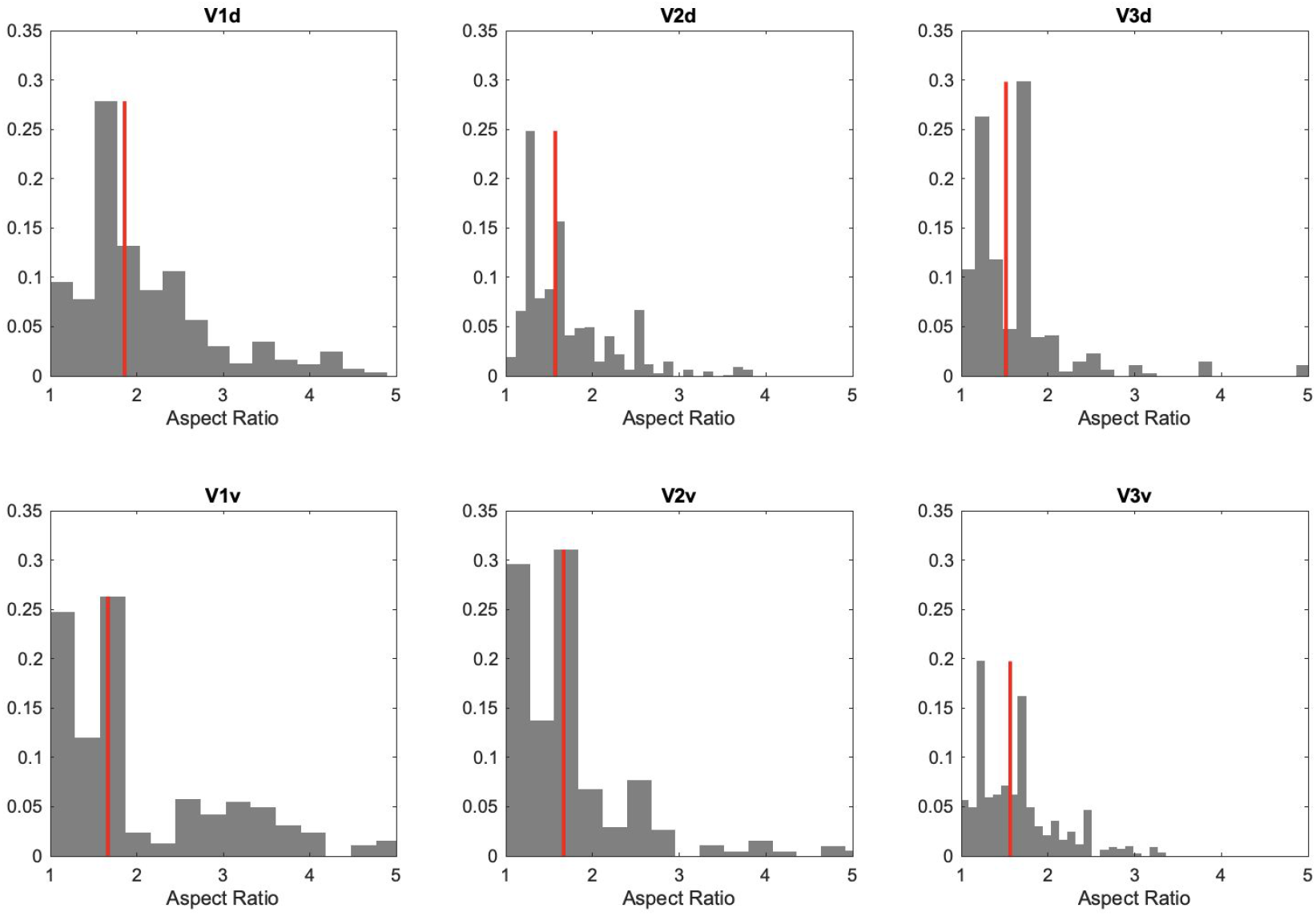
mrVista-elliptical. Experimental data. Aspect ratio histograms in dorsal and ventral V1-V3. See main text for the rest of details.

**Fig S8.**
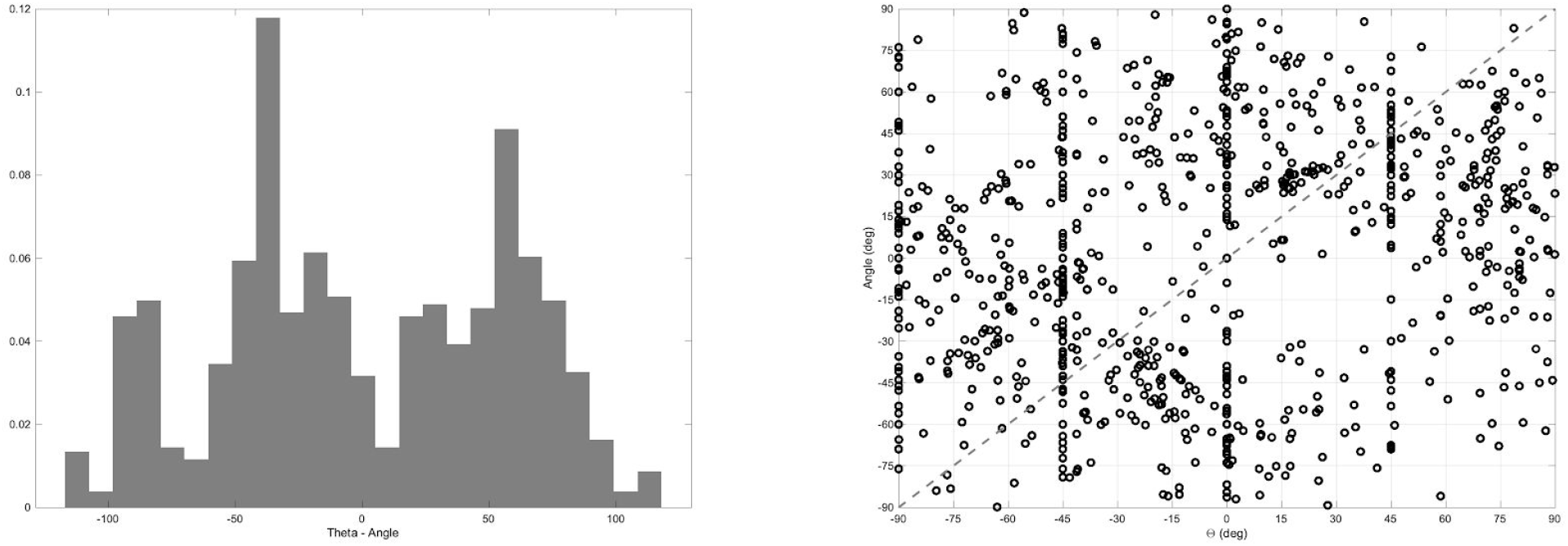
mrVista-elliptical. Experimental data. Comparison between the angle of the center of the pRF and the orientation Θ of the ellipse. (Left) Histogram of the difference between the angle of the pRF center and the orientation of the ellipse. A distribution around 0 would mean radiality, an orientation toward the fovea. (Right) Scatterplot of the pRF angle and the orientation of the ellipse. See main text for the rest of details.

**Fig S9.**
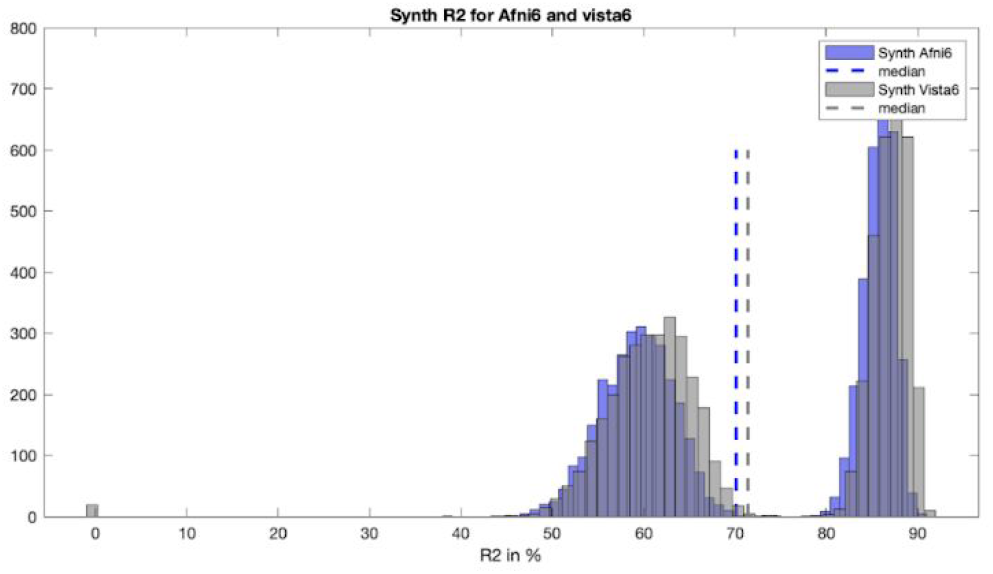
No bias in Vista-elliptical and AFNI-elliptical analyses of synthetic data. Histogram of the variance explained (R^2^) model fits to the synthetic data (300 sec, TR=1, ground truth aspect ratio = 1) analyzed with Vista-elliptical (gray) and AFNI-elliptical (blue) data. The thin dashed vertical lines represent the median values. The bimodal distribution of R^2^ values is explained by the two noise levels that were simulated, low and mid. The 95% confidence interval of the variance explained differences is [1.06-1.32] %, showing that practically there is no bias in favour of mrVista in how the data is synthesized.

**Fig S10.**
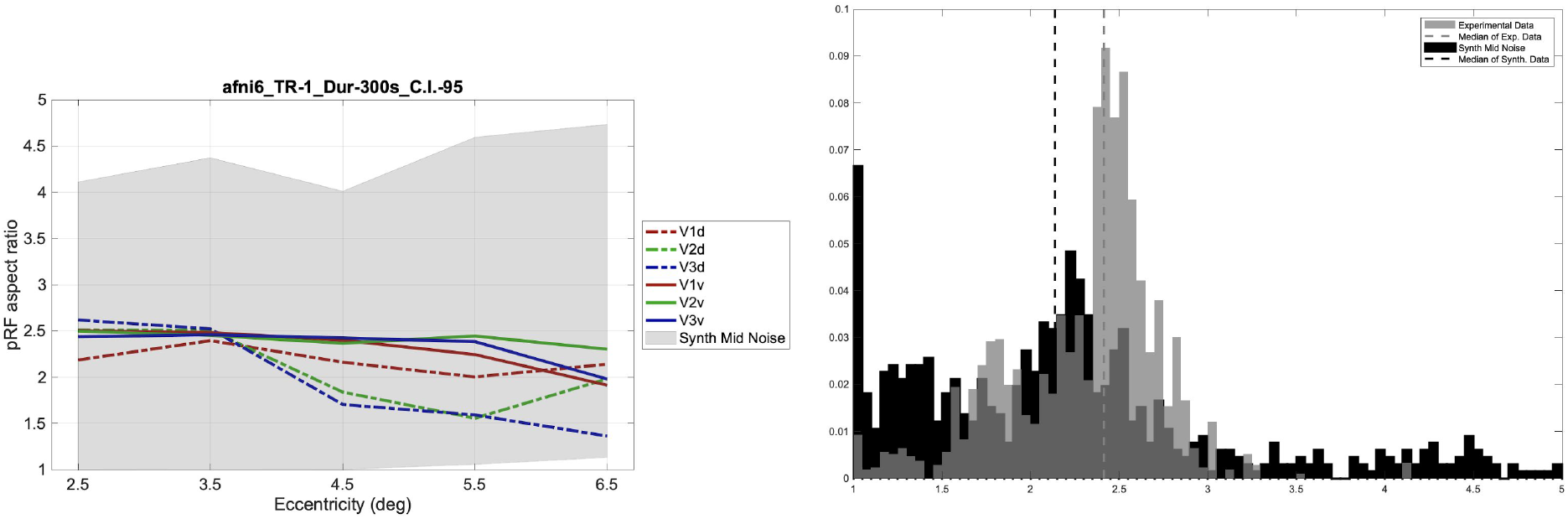
Experimental and synthetic data aspect ratio estimated with AFNI-elliptical. (Left) Estimated median pRF aspect ratios of experimental (color) data plotted as a function of eccentricity. The experimental data are plotted separately for ventral and dorsal regions of V1-3. Synthetic data were created using mid-level noise and is represented as a light gray band containing the central 95% aspect ratio values. (Right) Histograms of the estimated pRF aspect ratio for experimental (gray) and synthetic (black) data. Estimates were included in the histograms if the model fit explained at least 25 percent of the variance and the pRF position was between 2.5-6.5 deg and the pRF area size estimate was between 6.5-30 deg^2^. The ground truth aspect ratio for the synthetic data was 1. The thin dashed vertical lines represent the median values for the experimental and synthetic analyses.

